# Perfusable 3D models of ureteric bud and collecting duct tubules

**DOI:** 10.1101/2025.06.19.659798

**Authors:** Kayla J. Wolf, Ronald C. van Gaal, Sebastien G.M. Uzel, Jonathan E. Rubins, Aline N. Klaus, Amelie Printz, Pooja Nair, Katharina T. Kroll, Paul Stankey, Lisa M. Satlin, Jennifer A. Lewis

## Abstract

Recent protocols have emerged to derive ureteric bud (UB) and collecting duct (CD) organoids directly from human induced pluripotent stem cells (hiPSCs). However, these 3D kidney tissues lack biophysical cues from luminal flow and a drainage outlet. To address these limitations, we have created perfusable 3D models of UB and CD tubules. UB organoids are first generated from hiPSCs followed by their dissociation into individual UB cells. Individual UB cells are then seeded onto a 3D perfusable channel embedded within an extracellular matrix composed of fragmented basement membrane matrix and collagen I, where they self-assemble into a confluent monolayer. During *in vitro* perfusion, these cells exhibit UB-like marker expression over several weeks, during which they undergo budding akin to early branching morphogenesis in developing kidneys. To further promote network formation, UB cells are bioprinted adjacent to a perfusable UB tubule, which form interconnections through luminal fusion. Finally, these 3D perfusable UB tubules are differentiated into collecting duct tubules under luminal flow. Our platform facilitates fundamental understanding of human collecting duct formation during renal development, while paving the way for using these physiologically relevant models for drug testing, disease modeling, and, ultimately, integration into bioprinted kidney tissues for therapeutic use.

## Introduction

Patient-specific kidney tissues derived from human induced pluripotent stem cells (hiPSCs) are needed for applications ranging from “clinical trials in a dish”^1^ to renal replacement therapy.^2,3^ During renal development, the ureteric bud (UB) undergoes branching morphogenesis from the Wolffian duct forming a CD network that bridges nephrons with the ureter, allowing filtrate removal.^4^ However, to date, most efforts have focused on creating kidney tissues by generating metanephric mesenchyme (MM)-derived organoids. While the nephron-rich organoids do contain glomerular, proximal and distal tubular segments with cellular complexity akin to first-trimester kidney, they lack ureteric bud (UB)-derived collecting duct (CD) networks.^5–8^ To address this limitation, recent protocols have emerged for generating UB/CD organoids, which typically involve embedding single organoids in 3D matrices that contain blind-ended tubules.^9–15^ However, UB/CD organoids lack biophysical cues that arise from luminal flow as well as a common drainage outlet.^9,10,16–20^

Existing UB and CD tubule-on-chip models provide controlled platforms for perfusion, but neither UB branching morphogenesis or UB-to-CD differentiation has been demonstrated. Kimura *et al.* found that fluid flow across a 2D UB cell monolayer modestly influences UB phenotype; however, they did not investigate CD differentiation.^21^ In fact, most CD tubule-on-chip models reported to date are composed of mouse, rat, or canine cells, including our prior efforts to create 3D perfusable CD tubules embedded in matrix.^22–25^ Recently, Hong and Song, *et al.* developed a perfusable CD on-chip derived from human iPSCs by superfusing a 2D monolayer and showed that osmolarity, along with fluid flow to a lesser extent, enhances CD maturation.^26^ However, their model relied on 2D monolayers grown on rigid, non-degradable surfaces, which prevent CD cells from forming 3D lumens or branching architectures. While collectively these models provide some mechanistic insight, they are incapable of supporting branching morphogenesis into 3D ureteric bud or collecting duct networks.

Here, we report the biofabrication of a perfusable 3D UB tubule model that exhibits budding into the surrounding extracellular matrix, luminal fusion with bioprinted UB networks, and differentiation into a 3D CD tubule on chip. These 3D tubule models are produced by first differentiating hiPSCs to form UB organoids and then dissociating into individual UB cells. Next, these organoid-derived UB cells (oUBs) are seeded onto a cylindrical channel embedded in an extracellular matrix composed of collagen I (Col I) and basement membrane matrix (BsM), which supports UB monolayer formation, budding, and embedded bioprinting. Our matrix is specifically designed to achieve these key attributes by suspending granular BsM fragments in a fibrous Col I network. When oUBs cells are seeded onto this matrix, they form a 3D tubular monolayer that exhibits both budding and branching. We further demonstrate that UB networks, patterned adjacent to these perfusable tubules via embedded bioprinting, form luminal connections. Through further differentiation under luminal flow, we transformed these perfusable UB tubules to CD tubules on chip. Our work opens new avenues for creating physiologically relevant, 3D kidney tissues for drug testing, disease modeling, and therapeutic use.

## Results

### I. Matrix optimization

We initially focused on developing an extracellular matrix that (i) supports monolayer formation by UB cells in matrix encapsulated channels, and (ii) allows for UB branching morphogenesis within the matrix to form interconnected networks (**Fig. 1A**). UB organoids are generated from hiPSCs by adapting the protocol reported by Zeng and Huang *et al.*^9^ Unlike other methods, this protocol yields highly expandable UB organoids enriched with UB tip-like cells.^10,11^ These UB organoids are dissociated to yield oUBs, which are cultured in 2D for 2 days on four candidate matrices (**Fig. 1A**). While gelatin-fibrin (gelbrin) matrices support confluent monolayers composed of immortalized and organoid-derived proximal tubule cells^27,28^, we find that few oUBs adhere to this matrix, and those cells that did exhibit a mesenchymal-like morphology (**Fig. 1B-C**). By contrast, basement membrane matrix (BsM) is widely used for UB organoid differentiation.^9,10^ Both Matrigel and Geltrex contain laminins and collagen IV, which are known to promote renal epithelial cell adhesion and are expressed in renal tubular basement membranes.^29–31^ Interestingly, oUBs seeded on a 50% (vol/vol) BsM do not form confluent monolayers, while those seeded on 1 mg/mL Col I matrices form a monolayer that lacks a clear epithelial-like cobblestone morphology (**Fig. 1B**). However, when these cells are seeded on a hybrid matrix composed of 50% (vol/vol) BsM and 1 mg/mL collagen type I (Col I), they achieve nearly 100% confluency after 2 days (**Fig. 1C**). These monolayers exhibit the characteristic cobblestone morphology expected for epithelium. Hence, this BsM - Col I matrix is optimal for supporting epithelial monolayer formation.

**Figure 1.**
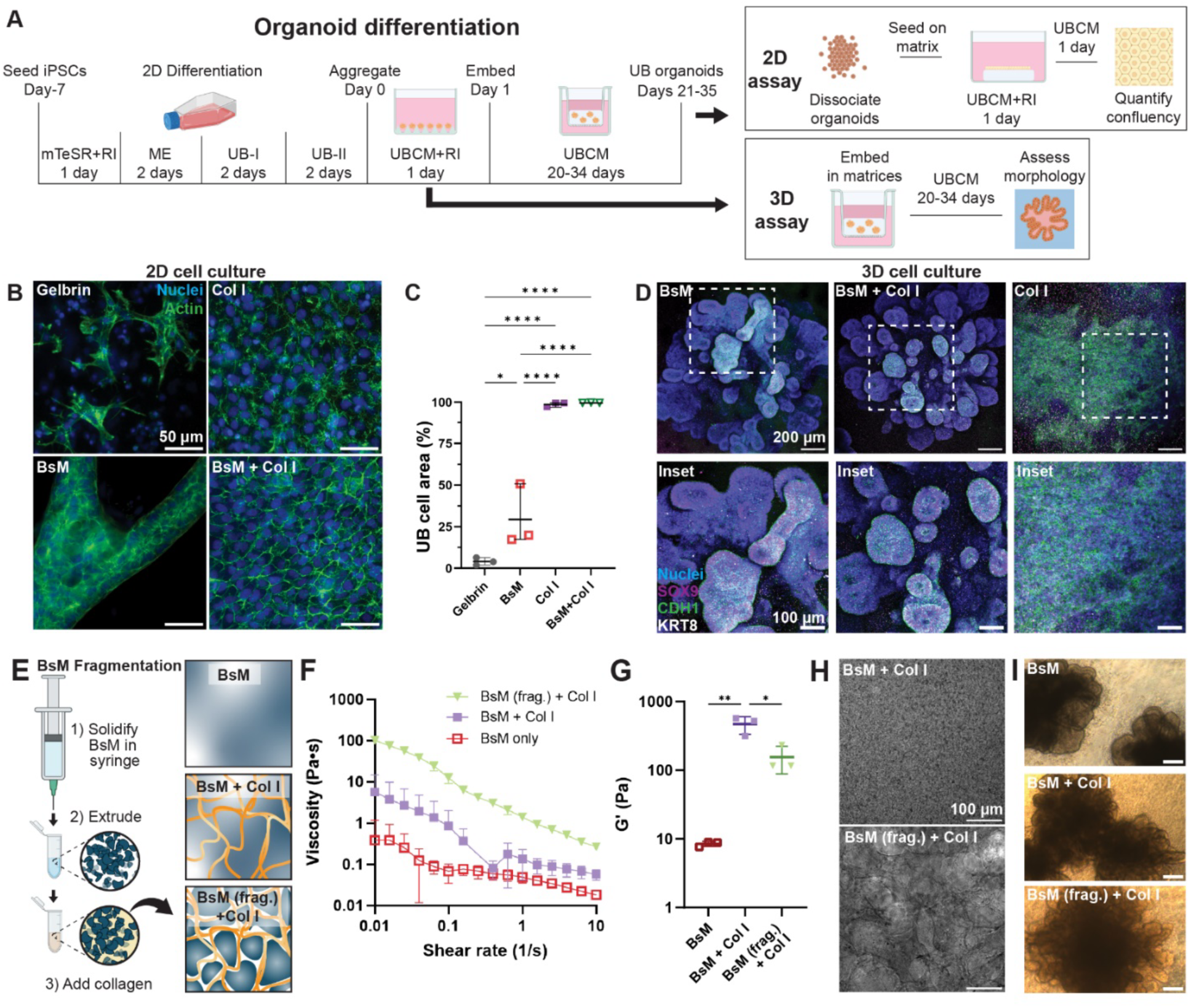
Matrix optimization. **A**) Differentiation protocol (adapted from Zeng & Huang *et al*.^9^) and subsequent use of UPCs for 3D budding assay and oUBs for 2D cell adhesion assay and on chip culture. **B**) Immunofluorescence images of 2D cell adhesion assay, in which cells are stained for actin cytoskeleton (green) and nucleus (blue) on matrices (gelbrin, 50% vol/vol BsM, 1 mg/mL Col I, and 50% vol/vol BsM – 1 mg/mLCol I) after two days of culture on matrix. Scale bar = 50 µm. **C**) Confluency quantification of the oUB monolayer on different matrices. Data points represent N=3 independent experiments (different cell and matrix batches). Statistical analysis by one-way ANOVA followed by Tukey’s multiple comparisons test. **D**) Immunofluorescence images from 3D budding assay of UB organoids in 50% vol/vol BsM, 50% vol/vol BsM-1 mg/mL Col I and 1 mg/mL Col I matrices (from left to right) and region of interest (inset; bottom row). Stained for nucleus (blue), SOX9 (red), CDH1 (green), KRT8 (white) at day 21 of differentiation. Scale bar = 200 µm, inset = 100 µm. **E**) Schematic overview of fragmented matrix preparation (left), and matrix formulations (50% vol/vol BsM, 60% vol/vol BsM – 1.5 mg/mL Col I, 60% vol/vol BsM (frag.) – 1.5 mg/mL Col I; right). **F**) Flow sweeps at 4 °C before collagen crosslinking and **G**) Storage modulus after collagen crosslinking of BsM, BsM - Col I, BsM (frag.) - Col I using the same formulations from subpanel E. Data points represent N=3 independent matrix batches. Error bars denote mean and standard deviation. **H**) Bright field imaging of matrices stained with picrosirius red comparing Col I network in BsM – Col I and BsM (frag.) – Col I matrices with same matrix composition as subpanel E. Scale bar = 100 µm. **I**) Phase images of day 21 UB organoids cultured in matrix formulations. Scale bar = 200 µm. Throughout the figure: *p<0.05, **p<0.01, ***p<0.001, ****p<0.0001.

Next, we explored the potential of each candidate matrix to support 3D budding during UB organoid differentiation. We aggregated day 0 UB progenitor cells (UPCs) overnight and then embedded them in these different matrices, where they were cultured in UB complete medium (UBCM) for up to 21 days (**Fig. 1A**). These UPC aggregates rapidly degraded gelbrin over the course of 2-3 days preventing further analysis (**Fig. S1A**). By contrast, multicellular aggregates embedded in Col I expressed canonical UB markers (e.g., SOX9, CDH1 and KRT8) after 21 days; however, these cells failed to form epithelial lumens (**Fig 1D, S1B**). We find that UPC aggregates embedded in either pure BsM or BsM-Col I exhibited budding with epithelial lumens expressing SOX9, CDH1 and KRT8 during organoid differentiation and growth (**Fig. 1D, S1C-E**). Our findings are consistent with prior studies that find epithelial cells embedded in BsM rapidly form luminal structures.^32^ Importantly, for the four matrices tested, only BsM-Col I matrix supports both UB monolayer formation and budding.

Since *in vitro* UB models have limited capacity to branch and elongate^9,10,16^, we further optimized our BsM - Col I matrix for embedded bioprinting. Granular hydrogels are known to exhibit both the requisite viscoelasticity and self-healing behavior for embedded bioprinting.^33–36^ To create BsM fragments, we solidified 100% (vol/vol) BsM in a syringe and then extruded this dense matrix through a 32-gauge nozzle (**Fig. 1E**) to create fragmented particles with cross-sectional areas ranging from ∼100 µm^2^ to 10,000 µm^2^ (**Fig. S2A-B**). By mixing these BsM particles with cold collagen (4 °C), we formed a heterogenous matrix composed of 60% (vol/vol) BsM fragments suspended in 1.5 mg/mL Col I. This mixture can be maintained for >2 h at this temperature with minimal collagen crosslinking resulting in a narrow, yet suitable, printing window. At 4 °C the matrix exhibited a shear elasticity of ∼3 Pa, a shear yield stress of ∼1 Pa and a 10-fold higher apparent shear viscosity (∼100 Pa·s at 0.01 Hz) compared to either non-fragmented 60% vol/vol BsM-1.5 mg/mL Col I or 50% vol/vol BsM controls. Importantly, neither control matrix exhibited a measurable shear yield stress (**Fig. 1F, Fig. S2C**).

Upon incubation at 37 °C, Col I forms a continuous fibrous network that surrounds the BsM fragments during matrix solidification. The storage modulus of 60% vol/vol BsM (frag.) - 1.5 mg/mL Col I (∼160 Pa) is significantly lower than non-fragment 60% vol/vol BsM – 1.5 mg/mL Col I (∼470 Pa), yet still an order of magnitude more than 50% vol/vol BsM alone (∼10 Pa) (**Fig. 1G, Fig. S2D**). Staining with picosirius red revealed collagen fiber bundles that are qualitatively longer in BsM (frag.) - Col I matrices compared to their non-fragmented counterparts (**Fig. 1H**), which correlates with lower crosslinking density and stiffness.^37^ Finally, we confirmed that BsM fragmentation did not negatively impact UB phenotype during differentiation. Phase imaging revealed that both UB organoid size and morphology are similar in 50% vol/vol BsM, 60% vol/vol BsM – 1.5 mg/mL Col I, and 60% vol/vol BsM (frag.) – 1.5 mg/mL Col I matrices (**Fig. 1I**). Transcriptomic analysis of 172 genes consisting of UB, CD, and off-target markers revealed no significant differences in organoids cultured in each of these matrices (**Fig. S3**). Hence, the 60% vol/vol BsM (frag.) – 1.5 mg/mL Col I matrix supports monolayer formation, branching morphogenesis, and embedded bioprinting. This optimized matrix, referred to as BsM (frag.) – Col I, is used for all further studies.

### II. 3D UB tubule fabrication

We used a pin pull-out approach to fabricate our perfusable 3D tubules models.^22,28^ Our modular chip platform provides easy access to the basal compartment for media changes, enables static and perfused tubule culture, and facilitates embedded printing directly into the surrounding matrix (**Fig. 2A-B, S4A-B**). The oUB cells are seeded at ∼100 x 10^6^ cells/mL in each cylindrical channel, where they assemble into a confluent monolayer that buds into the matrix by ∼1 week after seeding (**Fig. 2C**). A difference in the medium height (and, hence, pressure) between the apical and basal reservoirs is sufficient to change the tubule diameter encapsulated within BsM (frag.) - Col I matrices (**Fig. S4C**). While this phenomenon could be exploited for inducing physiological stretch in tubules, it could be problematic if it gives rise to tubule collapse or bursting. We systematically varied the pressure difference between the apical and basal compartments and measured the corresponding channel diameter in absence of epithelium (**Fig. S4C**). The channel diameter changes from its original value of 1 mm to values ranging between ∼0.450 mm to ∼1.1 mm depending on medium height. To maintain a tubule diameter of ∼1 mm, we kept the medium volume the same in each chamber, yet we located the apical reservoir at a slightly higher height during medium changes to avoid tubule collapse. Notably, the presence of BsM (frag.) - Col I leads to more stable tubule dimensions compared to non-fragmented BsM – Col I controls (**Fig. S4C**). The tubules can be perfused at volumetric flow rates ranging from 5 µL/min to 66 µL/min for >1 week without monolayer disruption despite changes in their cross-sectional geometry (**Fig. 2C**).

**Figure 2.**
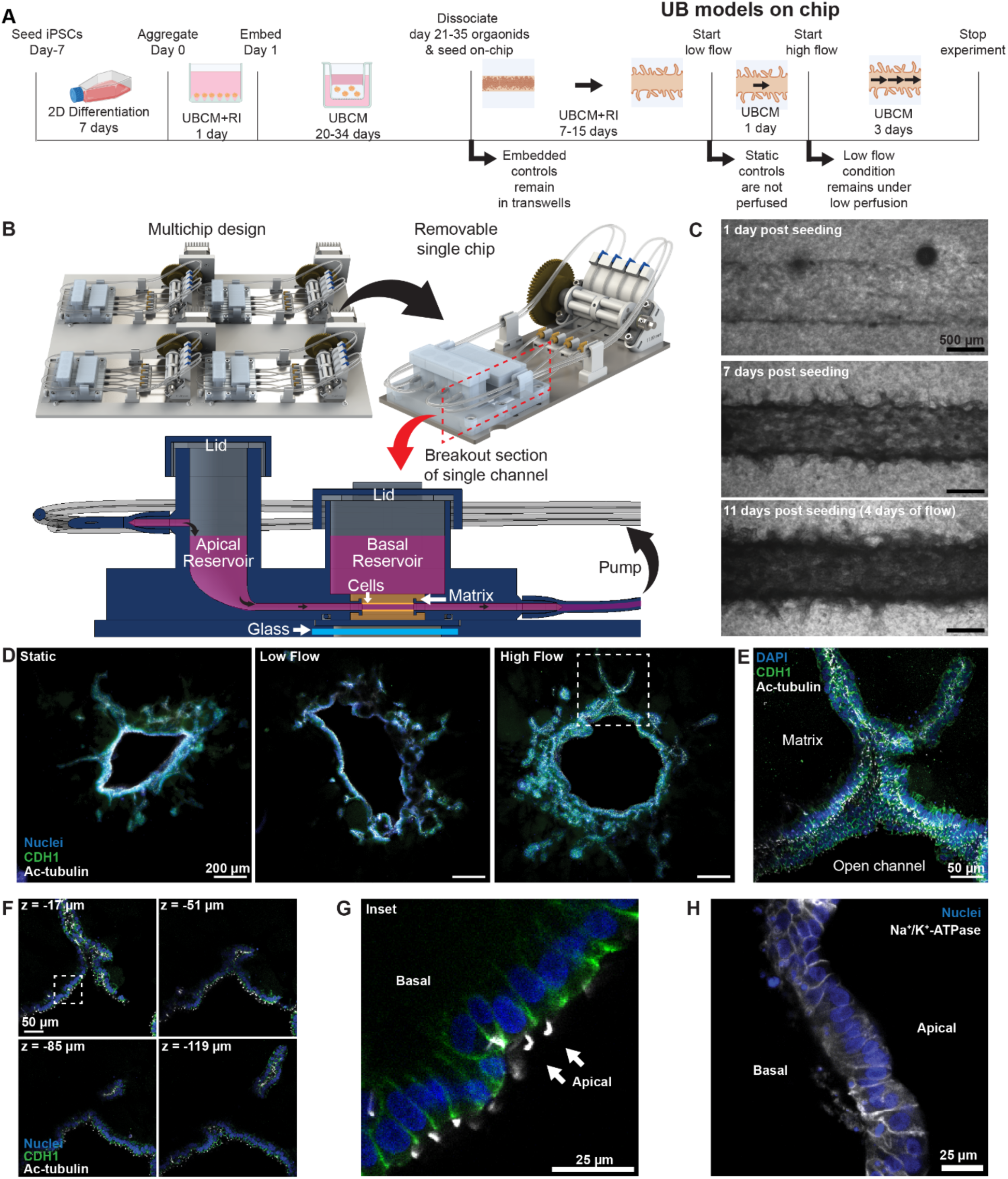
Perfusable 3D UB tubules on chip. **A**) Experimental timeline for UB organoid differentiation, static on chip culture, and luminal perfusion. **B**) Schematic overview of modular chips with integrated pumps (top) and cross section of a single channel within a chip (bottom) highlighting tubule culture separating apical and basal compartments. **C**) Phase images of oUB cultured for 11 days on chip. Scale bar = 500 µm. **D**) Immunofluorescence images of 3D UB tubule cross-sections from static, LF, and HF conditions stained for nucleus (blue), cell-cell junctions via CDH1 (green), and cilia via Ac-tubulin (white). Scale bar = 200 µm. Dashed white box marks a region of interest with protruding bud. **E**) Maximum intensity projection of a protruding bud in the region of interest from subpanel B and **F**) montage of single z-slices of same region of interest at different depths. Scale bar = 50 µm. **G**) Inset shows monolayer with apically oriented single cilia (arrows). Scale = 25 µm. **H**) Representative immunofluorescence image of Na+/K+-ATPase (white) expression and nuclei (blue) in cross-section of UB tubule perfused under HF. Scale bar = 25 µm.

Next, we characterized bud morphology to determine cell polarity and lumen connectivity with central UB tubules (∼ 1 mm in diameter) under both low (LF; 5 µL/min) and high (HF; 66 µL/min) volumetric flow rates (**Fig. 2D),** which correspond to fluidic shear stress values of 0.006 dyn/cm^2^ and 0.08 dyn/cm^2^, respectively. After perfusing these 3D UB tubules for 4 days, they are fixed, stained and imaged using confocal microscopy. Cross-sectional images of the 3D tubules reveal the presence of a confluent monolayer that is connected to UB buds that protrude into the matrix (**Fig. 2D-E, Fig. S5A**). Individual z-slices of a bifurcated branching tubule reveal an invagination in the central lumen that is continuous with cilia-positive apical cell surfaces (**Fig. 2E-F, Fig. S5B**). CDH1^+^ cell-cell junctions are observed with apically oriented cilia in all conditions (**Fig. 2D-G**). Importantly, budding lumens are continuous with the central perfusable tubule (**Fig. 2E-F**). UB tubules exhibit normal barrier function, as evidenced by retention of FITC-inulin within the perfused channel relative to non-epithelialized controls (**Fig. S6A-C**). Each set of perfusable 3D UB tubules is generated using dissociated oUB cells obtained from different UB organoid batches. In all cases, UB bud formation and protrusion into the matrix is observed, albeit with some variation in budding number and density (**Fig. S6D-E**). Despite such differences, the maximum bud length, which ranges between roughly 200-600 µm did not vary significantly (**Fig. S6G**). Finally, these 3D UB tubules exhibit Na+/K+-ATPase transporter expression, which is ubiquitously expressed on their basolateral surface, as required for sodium homeostasis (**Fig. 2H**).^38^

### III. Bioprinting UB tubule networks

To investigate whether bioprinted UPCs will self-assemble into tubules that undergo fusion, we printed such cells in a sterile tissue culture hood using a low-cost, open-source printer recently reported by Weiss and Mermin-Bunnell *et al.*^39^ equipped with a custom, chilled build plate (**Fig. S7A-B**). Printed UPCs consistently expanded and differentiated into budding structures during culture in UBCM over 21 days after printing (**Fig. S7C-G**). Using embedded bioprinting, we patterned alternating filamentary features composed of UPCs that express either GFP or mCherry within BsM (frag.) – Col I and tracked these two cell populations over 21 days. Within the first 7 days, the GFP- and mCherry-expressing UPC populations began to form tubules that undergo fusion with one another and result in contiguous lumens by day 21 (**Fig. S7H**). Immunofluorescence and confocal imaging confirmed bud-to-bud connections form between these two cell populations (**Fig. S7I**).

To create a physiologically relevant architecture, we printed a branching UPC network adjacent to a 3D channel seeded with UPCs followed by culture on chip for 7 days (**Fig. 3A**). Note, UPCs are used for embedded bioprinting, because they can be made into a bioink immediately after 2D culture. To induce rapid budding within this 7-day period (i.e., rather than the typical 21-day differentiation timeline), the printed UPCs are cultured in UB differentiation media following the protocol reported by Shi and McCracken *et al.*^10^ (**Fig. 3B**). Importantly, we find that the printed branching networks exhibit additional budding of smaller UB branches that protrude into the surrounding matrix yielding a hierarchically branching network (**Fig. 3B)**. Surprisingly, the UPCs seeded within the central 3D tubule did not form a confluent monolayer even though they did undergo budding (**Fig. 3B**).

**Figure 3.**
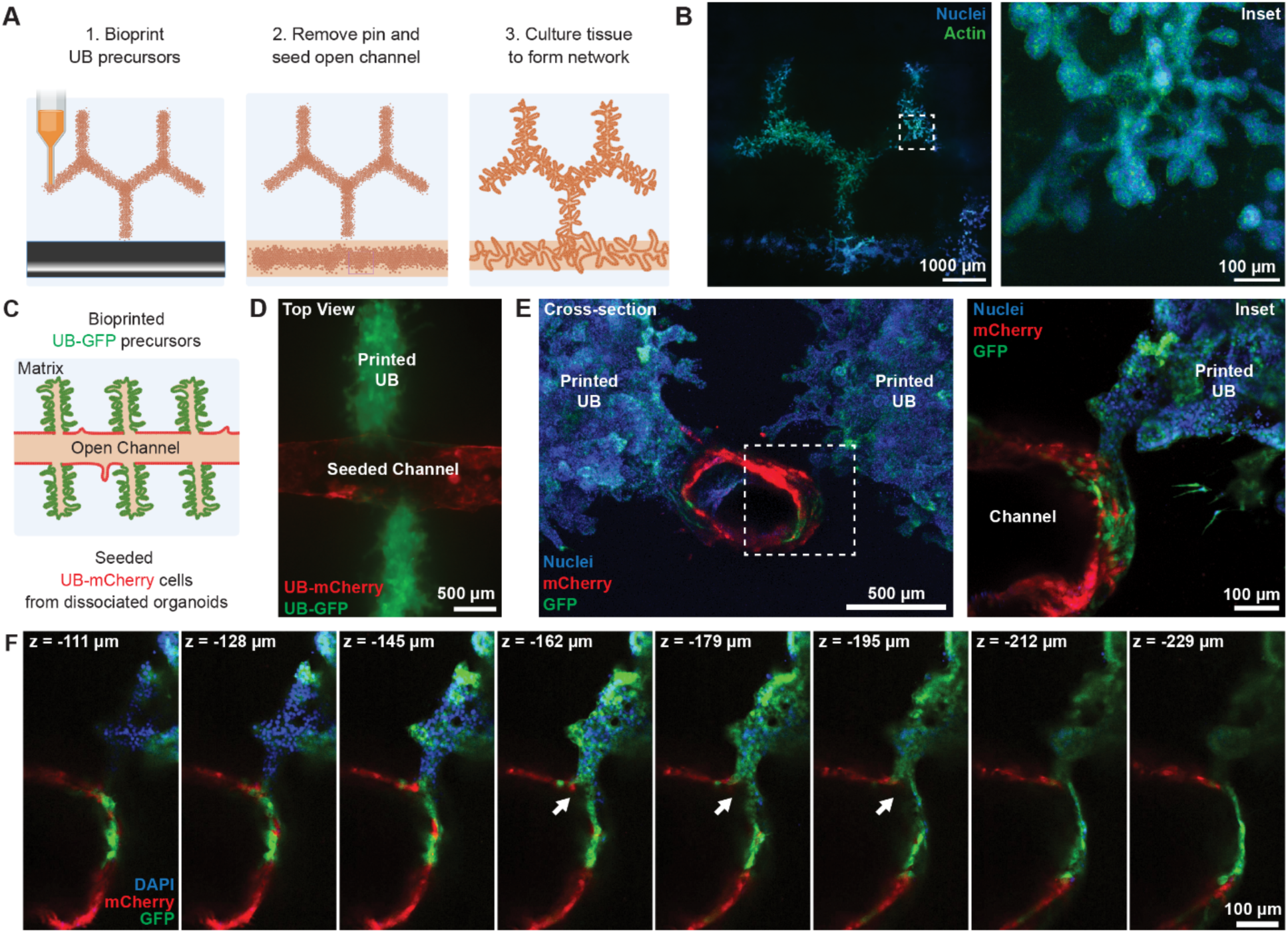
Branching UB networks via embedded bioprinting. **A**) Schematic overview of complex branching network produced by embedded bioprinting a UPC bioink adjacent to a central lumen produced by pin pullout. **B**) Immunofluorescence images of printed UPC network and seeded channel (left), where the dashed box indicates region of interest (higher magnification; right). Cells were cultured for 7 days after printing and fixed/stained for actin (green) and nuclei (blue). Scale bar = 1000 µm, inset scale bar = 100 µm. **C**) Schematic view of multiple UPC filament printed adjacent to central UB tubule. Lineage tracing with GFP- and mCherry-expressing cells delineates printed and seeded tubule cell populations. **D**) Epifluorescence image of printed UB-GFP cells fusing with UB-mCherry cells seeded in central tubules (image taken on day 25 after printing; top view). Scale bar = 500 µm. **E**) Maximum intensity projection of cross-section (image taken on day 32 after printing) (left) where dashed line indicates region of interest (higher magnification; right) showing nuclei (blue), GFP (green), and mCherry (red). After fixation, sample was treated with easy index. Scale bar = 500 µm, inset scale bar = 100 µm. **F**) Single z-slices of region of interest in subpanel E indicating tubule connection and cellular migration from printed branch to main channel. White arrow indicates invagination resulting from lumen-to-lumen connection. Scale bar = 100 µm.

We next focused on driving connections between printed UB features and perfusable 3D tubules by seeded the cental lumen with oUBs, since these cells readily form confluent monolayers (**Fig. 3C**). For each chip, we printed 6 UPC-laden filaments (4 mm in length) adjacent to central UB tubule such that there were at least 6 locations in which lumen-on-lumen connections could potentially form (**Fig. 3C**). After the UPCs features are printed and oUBs are seeded into the central channel, the constructs are cultured for ∼21 days or longer to enable UPC differentiation and the formation of a confluent oUB epithelium in the central 3D tubule (**Fig. 3D**). Importantly, we find that fusion of bioprinted tubules and the central UB tubule occurs, which results in lumen-to-lumen connections between UPC and oUB cell populations (**Fig. 3E-F**).

### IV. 3D UB-to CD tubule differentiation on chip

To promote the desired UB-to-CD tubule differentiation, we first generated 3D UB tubules composed of oUB monolayers following our 7-day protocol followed by the introduction of a medium that contained aldosterone and vasopressin, but not UB-promoting factors (**Fig. 4A; Table S3**). Note, this medium has been used for further differentiating UB organoids to CD organoids in static culture.^9^ We find that 4-6 days of UB organoid culture in CD medium is sufficient to both decrease UB marker expression (e.g., RET) and increase CD marker expression (e.g., WNT9B) resulting in principal-like CD cells.^9,10^ To evaluate flow effects on their differentiation, we compared CD and UB tubules subjected to LF and HF luminal perfusion to two controls: (1) whole organoids embedded in the same BsM (frag.) - Col I matrix and (2) 3D tubules differentiated under static (no flow) conditions on chip (**Fig. S8**).

**Figure 4.**
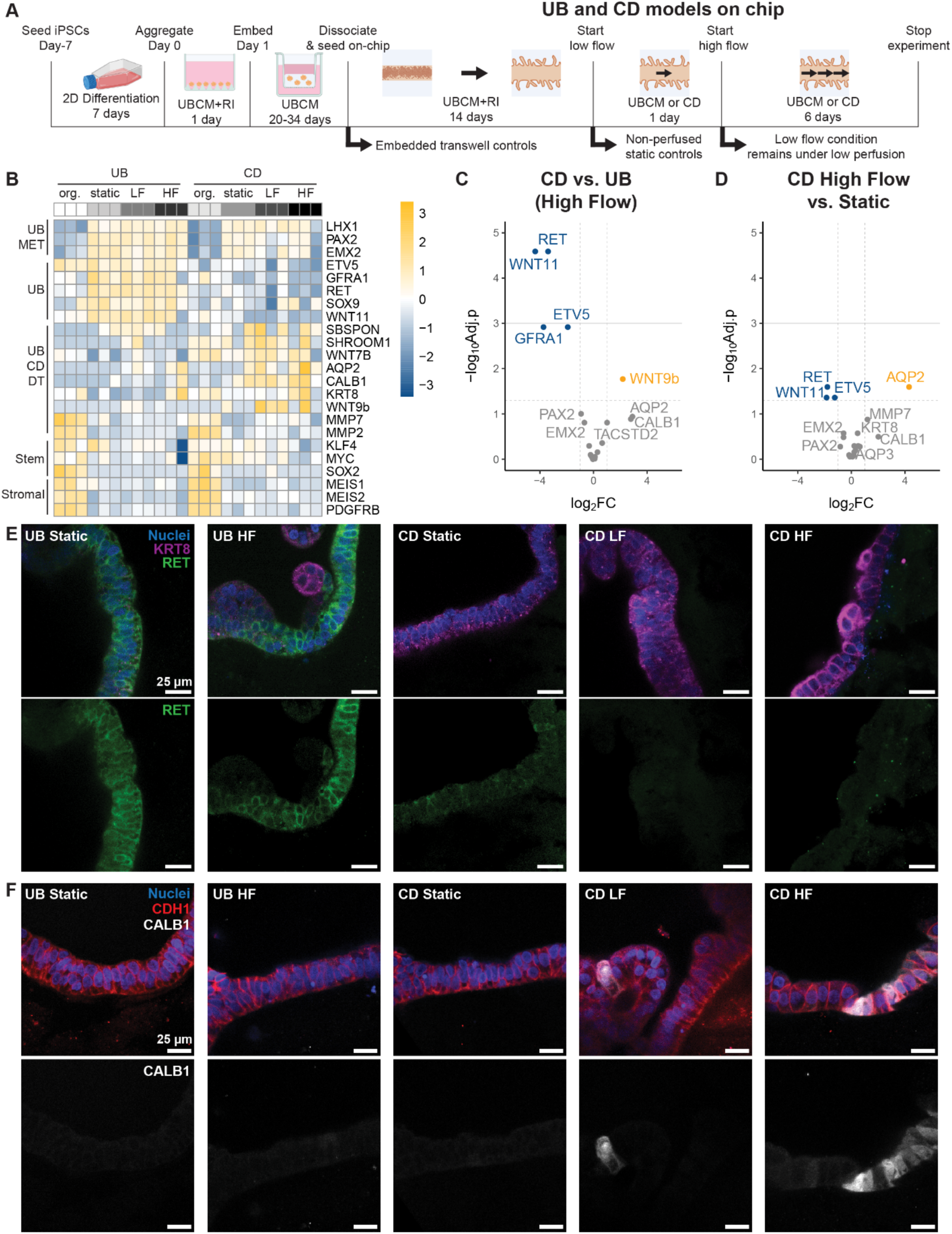
3D UB-to-CD tubule differentiation under perfusive flow. **A**) Experimental timeline including 7-day CD differentiation concurrent with perfusion. **B**) Heatmap of mRNA expression of selected genes from UB and CD embedded whole organoids (org.) and tubules on chip in static, LF, and HF conditions. Columns in each condition represent N=3 independent experiments. Colors correspond to up- (orange) and down- (blue) regulated genes scaled to the standardized z-score. MET (mesenchymal to epithelial transition), DT (distal tubule), Stem (stemness markers). Volcano plots depicting differential mRNA expression of UB and CD markers in **C**) HF CD on chip relative to HF UB on chip and **D**) HF CD on chip relative to static CD on chip. Colors correspond to non-significantly expressed genes (gray), upregulated genes (orange), and downregulated genes (blue). Dashed horizontal line = adjusted p<0.05, solid horizontal line = adjusted p<0.01, vertical line = 2-fold change. Data points represent individual genes from N=3 independent experiments where p-values are adjusted by the Benjamini-Hochberg correction. **E**) Representative immunofluorescence images of markers in all chip conditions. Top row: nuclei (blue), KRT8 (magenta), RET (green). Bottom row: RET (green). Scale bar = 25 µm. **F**) Representative immunofluorescence images of markers under in all chip conditions. Top row: nucleus (blue), CDH1 (red), CALB1 (white). Bottom row: CALB1 (white). Scale bar = 25 µm.

We first assessed gene expression of UB, CD, and off-target markers using a 48-gene panel. Notably, whole UB and CD organoids expressed more stromal markers (e.g. *MEIS1/2*, *PDGFRB*) and more stemness markers (e.g. *KLF4*, *MYC*, *SOX2*) than static or perfused 3D UB and CD tubules on chip (**Fig 4B, S9, S10**). By contrast, UB and CD markers (e.g. *PAX2*, *RET*, *EMX2*, *WNT9b*, *WNT11*) are upregulated on chip under both static and flow conditions relative to embedded organoid controls. Together, these findings indicate that the UB/CD-like gene expression of each cell population is enriched merely by seeding them as 3D tubules on chip. The presence of stromal and undifferentiated cells is likely reduced during organoid dissociation, since these cells are more likely to invade and adhere to BsM that is broken up and discarded. Notably, whole organoids embedded in BsM, BsM-Col I, and BsM (frag)-Col I exhibit no significant differences in gene expression with respect to matrix composition (**Fig. S3**). Transcriptomic analysis also revealed that both UB organoid (controls) and 3D UB tubules differentiated to CD organoids and tubules, respectively, exhibited increased CD marker expression (e.g., WNT9B) and decreased UB marker expression (e.g., *WNT11*, *RET*, *ETV5*, *GFRA1*) relative to UB controls across multiple flow conditions. Hence, we can successfully differentiate 3D UB tubules to 3D CD tubules on chip (**Fig. 4B-C, S9, S11A-C**).

Since these 3D tubules contract during monolayer formation, there are variations in tubule diameter and, hence, shear stress experienced by these confluent cell monolayers at a given volumetric flow rate (**Fig. S8**). We estimate that maximum fluidic shear stresses at LF (average: 0.031±0.015 dyn/cm^2^) and HF (average: 0.34±0.14 dyn/cm^2^; **Fig. S8**) deviate by up to an order of magnitude from the applied fluidic shear stresses of 0.006 dyn/cm^2^(LF) and 0.08 dyn/cm^2^ (HF) calculated based on their initial tubule diameter (1 mm). Despite this variability, CD tubules subjected to HF exhibited a reduction in *RET*, *WNT11*, and *ETV5* expression, but a higher *AQP2* expression compared to static tubule controls (**Fig. 4D**, **S11D-F**). By comparison UB marker expression is not affected in the same manner apart from a significant upregulation of *UPK2* and *AQP2* under HF compared to static UB tubule controls (**Fig. S11G-I).**

Using immunofluorescence and confocal imaging, we further investigated RET expression in both 3D UB and CD tubules, since RET is a receptor for GDNF and a canonical marker of UB tip cells. In good agreement with our transcriptomic analysis, we observed qualitatively less membrane-localized RET expression in CD compared to UB tubules, particularly for tubules subjected to flow (**Fig. 4E, S12**). Downregulation of RET in CD tubules suggests that the UB to CD differentiation is likely enhanced under flow. We next investigated AQP2 expression, a critical water transport protein and canonical marker of principal cells in the collecting duct. In contrast to gene expression data, upregulation of AQP2 expression under flow is not observed (**Fig. S13**). Furthermore, we did not observe clear trends in membrane localization or polarization of AQP2 with respect to differentiation state or luminal flow rate (**Fig. S13**). It is possible that while flow affects AQP2 gene expression, protein expression and membrane trafficking are determined by other factors, such as stretch and osmolarity, which are not controlled or investigated here. Finally, we investigated the canonical CD marker CALB1. Although *CALB1* gene expression did not vary significantly between flow rates and differentiation states in this study, we observed heterogenous, higher CALB1 protein expression in cells exposed to flow compared to static controls, namely in CD and not UB (**Fig 4F, S14**). Collectively, our findings demonstrate that (i) we can differentiate UB to CD on chip under perfusive flow, (ii) our platform can be used to model CD development, and (iii) luminal flow may promote CD maturation.

## Discussion

We have addressed the critical need for 3D UB and CD tubule models derived from human induced pluripotent stem cells. Unlike recently reported 2D UB on chip models^21,26^, our model provides the necessary matrix microenvironment to support both branching morphogenesis and luminal perfusion. Additionally, we show that bioprinting into granular matter offers a powerful strategy to bypass the extended developmental timelines and limited branching often encountered *in vitro*^9,10,16^ enabling one to deterministically pattern and form branching network architectures. We further show that luminal flow can influence and potentially enhance UB to CD differentiation and maturation. Transcriptomic analysis and immunofluorescence staining suggest that luminal flow may reduce UB marker expression (especially RET, but also WNT11, GFRA1) and increase CD marker expression (e.g. AQP2, CALB1). Although it is not surprising that removal of GDNF from the medium during CD differentiation would lead to reduced RET expression, precisely how flow effects RET expression remains unknown. Inconsistencies between gene and protein expression data for AQP2 and CALB1 may reveal the need for further improvement to CD differentiation protocols or microenvironmental cues. Nevertheless, we show that cues from luminal flow (e.g., shear stress, pulsatility, circumferential stretch, torque on cilia) do promote tubule maturation *in vitro*. Future embodiments based on genetically engineered hiPSC-derived cells would enable one to investigate malformations in the ureter and collecting ducts^40^ as well as mutations underlying CD-specific diseases, such as nephrogenic diabetes insipidus.^41^

Despite these promising findings, our understanding of branching morphogenesis, CD differentiation, and flow-driven tubule maturation remain limited. Single-cell RNA sequencing or proteomic analysis could further illuminate potential pathways and cell subpopulations that are differentially expressed under each condition, while gain/loss of function assays could elucidate the role of candidate pathways. While lumen-to-lumen connections between bioprinted branching networks and central 3D tubules formed leading to continuous pathways for fluid drainage, further efforts are needed to create more complex, fully perfusable tubule architectures.

In summary, we created perfusable 3D UB and CD tubules that may be used for drug discovery, disease modeling, and developmental biology applications. We also ultimately envision that these initial steps towards patterning complex branching tubule architectures may open new avenues for building human kidney tissues for therapeutic use, especially when combined with the ability to co-print nephron-rich tissues. We posit that this approach could ultimately lead to CD tubule fusion with nephrons for filtrate generation and removal.

## Materials and methods

Comprehensive lists of all resources and reagents including source and vendor information are provided in **Tables S1-5**.

### Growth factor preparation

The following growth factors were reconstituted in Dulbecco’s Phosphate Buffered Saline with calcium and magnesium (DPBS^+/+^) containing 0.1% w/v bovine serum albumin (BSA): Fibroblast growth factor 2 (FGF2) at 1 mg/mL, FGF7 at 100 μg/mL, glial cell line-derived neurotrophic factor (GDNF) at 100 μg/mL, R-spondin 1 (Rspo1) at 100 μg/mL and epidermal growth factor (EGF) at 200 μg/mL. The following small molecules were reconstituted in dimethyl sulfoxide (DMSO): Y27632 at 10 mM, LDN-193189 at 1 mM, CHIR99021 at 20 mM, TTNPB at 10 mM, A83-01 at 20 mM, Janus kinase (JAK) inhibitor I at 10 mM, SB202190 at 20 mM, and aldosterone at 10 mM.

### hiPSC culture

Human iPSCs (hiPSC; BJFF.6, male, provided by Prof. Sanjay Jain at Washington University) were cultured for maintenance on plates coated with 1% Matrigel in DMEM/F12 and cultured in mTeSR Plus (mTeSR+) medium. iPSCs were passaged at 60-80% confluency with ReLeSR (according to manufacturer’s protocols) at split ratios of 1:8 - 1:20. Frozen cell stocks were preserved in 1:1 embryonic cell freezing medium and mTeSR+ medium with 10 μM Y27632. Normal karyotype of cells was confirmed by WiCell. At least every six months, a routine mycoplasma screening was performed using the Lonza Mycoplasma Detection kit. iPSCs were cultured no more than 15 passages after karyotyping.

### hiPSC reporter line generation

BJFF.6 iPSC lines expressing GFP or mCherry were generated following manufacturer’s protocols using the piggyBac transposon system. Custom plasmids expressing eGFP, mCherry, or hyPBase under the EF1A or CAG promoter with puromycin resistance were procured from Vectorbuilder (plasmid design available upon request). Briefly, iPSCs were split into a 6-well plate coated with 1% Matrigel in DMEM/F12 at a 1:10 ratio and cultured with mTeSR+ until cells reached 60-70% confluency. On the day of transfection, two mixtures were prepared for each well. First, a tube containing 125 μL of Opti-MEM and 10 μL of Lipofectamine Stem Cell Reagent, and second, a tube containing 125 μL of Opti-MEM with 2.5 μg of plasmid for insertion (either GFP or mCherry) and 0.5 μg of hyPBase plasmid. Then, 125 μL of each mixture were combined in a separate tube and left to complex for 10 minutes. Meanwhile, cell medium was aspirated and cells were rinsed with 1 mL of Opti-MEM with 10 μM Y27632. Then, 1 mL of Opti-MEM with 10 μM Y27632 was added to each well as well as 250 μL of complexed solution. Cells were incubated for four hours and then medium was exchanged with 1.25 mL of mTeSR+. Cells were incubated for 24 hours and medium was exchanged with a full culture volume of mTeSR+. Medium was exchanged daily until cells reached 70% confluency, and then 1 μg/mL puromycin was added for 2 days to select cells. Culture was continued in 500 ng/mL for continued selection and regular passaging protocols were followed.

### UB organoid differentiation and dissociation

Human iPSCs were differentiated towards UB organoids following the protocol of Zeng and Huang *et al.* with minor adaptations from established protocols.^9^ Briefly, iPSCs were plated on a 1% Matrigel pre-coated 12-well plate, or into a T75 coated flask (cultured with 1 mL per well in 12-well plates or 21 mL in a T75). The following day medium was replaced with pre-warmed ME-medium (Table S4). At day 3, the cells received pre-warmed UB-I medium (Table S4), and medium was refreshed the following day. At day 5, the UB-I medium was replaced with double culture volume of UB-II medium, and refreshed the following day (Table S4). On day 7, differentiated cells were washed with Dulbecco’s Phosphate Buffered

Saline without calcium or magnesium (DPBS^-/-^) and then dissociated with Accumax solution for 3-10 min at 37 °C. Cells were resuspended in UBCM (Table S4) with 10 μm Y27632 (UBCM-Y). Cells were either directly used for chip and printing experiments or 4,000-10,000 cells were seeded per microwell in an AggreWell™ 800 24-well plate (Aggrewell) according to manufacturer’s protocols. Aggregates were embedded in 50% (v/v) Geltrex matrix within a transwell system the following day. Post gelation, UBCM was added to the outer compartment and changed every 2-3 days during subsequent differentiation. Cells were cultured at 5% CO2 at 37 °C.

Embedded organoids were harvested from Transwells at days 20 to 34. To dissociate the embedded organoids, organoids and matrix from 3-6 Transwells in a 6-well plate were collected into a 15-mL conical tube with 1 mL of DPBS^+/+^ and pipetted ∼10x with a 1-mL pipette tip. Subsequently, volume was supplemented to 10 mL of DPBS^+/+^. The mixture was centrifuged at 300g for 5 min, yielding a distinguishable cell pellet, gel fraction, and supernatant. The supernatant and gel layer were slowly removed by aspiration, and the cell layer was then incubated with 10 mL of Accumax on an orbital shaker for 15 min at 37 °C. Cells were again centrifuged and the medium and gel fractions were removed and replaced with 1 mL of 0.25% trypsin for 15 min at 37 °C on an orbital shaker. After incubation, cells were briefly vortexed (∼2-3 seconds) and centrifuged at 100g for 3 min with maximum deceleration yielding a distinct cell pellet. Accumax solution was aspirated and cells were washed 1x in 1-2 mL UBCM-Y followed by resuspension in in 1-2 mL UBCM-Y. The final cell solution was passed through a 40-µm cell strainer and counted.

### Extracellular matrices

Gelbrin was prepared as a mixture of gelatin and fibrinogen. To prepare the extracellular matrix (ECM) components, a 15% (w/v) gelatin solution was formed by adding gelatin powder (Type A, 300 bloom from porcine skin) to a warm solution (70 °C) of DPBS^-/-^. The gelatin was stirred for 12 h at 70 °C, after which the pH was adjusted to 7.5 with 1 N sodium hydroxide (NaOH). The solution was sterile-filtered and stored at 4 °C. A fibrinogen solution (50 mg/ml) was made from lyophilized bovine blood plasma protein dissolved at 37 °C in sterile DPBS^-/-^. The solution was held at 37 °C without agitation for >45 min to allow complete dissolution. The transglutaminase solution (60 mg/mL) was made from lyophilized powder dissolved in DPBS^-/-^ and gently mixed for 20 s. The solution was then held at 37 °C for 20 min and sterile-filtered before use. A calcium chloride (CaCl2) stock solution (250 mM) was prepared from CaCl2 pellets dissolved in sterile water. For the preparation of stock solutions of thrombin, lyophilized thrombin was reconstituted at 500 U/mL in sterile water and stored at –20 °C. Thrombin aliquots were thawed immediately before use. Before the gelbrin was cast, several components were mixed in advance at appropriate concentrations, including 10 mg/mL fibrinogen, 2% (w/v) gelatin, 2.5 mM CaCl2, and 0.2% (v/v) transglutaminase. This solution was then equilibrated at 37 °C for 15-20 min before use to improve the optical clarity of the gelbrin. Next, the solution was rapidly mixed 250:1 with stock thrombin solution, resulting in a final thrombin concentration of 2 U/ml. Within 2 min at 37 °C, soluble fibrinogen cures to a fibrin gel. For this reason, the Gelbrin solution must be cast immediately onto the culture plate after being mixed with thrombin and incubated for >30 min before use.

Type I bovine telocollagen (Col I) was prepared at 1 mg/mL final concentration. For gel, 1 part of chilled collagen solution was mixed with 8 parts sterile water and 1 part 10x PBS. The pH of the mixture was adjusted to 7.2–7.6 using sterile 1 N NaOH and pH was monitored using pH paper. To prevent rapid gelation, the solution was maintained on ice. Finally, the solution was dispensed into culture plates and incubated at 37°C for 30-60 min to gelate.

Several bulk BsM-based matrices were prepared. While Matrigel and Geltrex (LDEV-Free Reduced Growth Factor Basement Membrane) were used interchangeably, Geltrex was used for most of this study. BsM matrices were prepared at concentrations of 50-100% vol/vol by mixing BsM with cold DPBS^+/+^ and maintaining the solution on ice until casting. To gelate, matrices were incubated at 37°C for 30-60 minutes. Similarly, mixtures of BsM and collagen were formed by varying Geltrex concentration between 40-86% (vol/vol) Geltrex, 0-40% (vol/vol) DPBS^+/+^, and 10-15% (vol/vol) collagen I stock solution (10.0 mg/mL) giving a final Col I concentration of 1.0 -1.5 mg/mL. For BsM-Col I matrices, the BsM and cold DPBS^+/+^ were first mixed by vortexing for ∼20-30 seconds. The solution was maintained on ice while the Col I stock was added. The solution was again vortexed and maintained on ice until gelation in an incubator at 37°C for 30-60 minutes.

Methacrylated hyaluronic acid (MeHA) matrices were prepared for comparison in fragmentation studies. Briefly, MeHA (Nanosoft Polymers, 50 kDa) was dissolved at 10% (wt/vol) in DBPS^+/+^ overnight at room temperature (RT) to form a stock solution. A 10 mg/mL stock solution of dithiothreitol (DTT) was prepared in DBPS^+/+^. A lithium phenyl-2,4,6-trimethylbenzoylphosphinate (LAP) photoinitiator stock was formed by dissolving LAP in deionized sterile water overnight at 4°C to form a 5% wt/vol solution. MeHA was mixed with DPBS^+/+^ to form ECM with final concentrations of 1% (wt/vol) MeHA, DTT at 1 mg/mL, and 0.05% wt/vol LAP. These solutions were cast and equilibrated overnight at 37°C in a humidified incubator enabling equilibration. The following day, MeHA matrices were photocrosslinked with UV light with an Omnicure S2000 for 120s on each side.

Fragmented matrices were prepared by adapting a protocol of Muir *et al..*^33^ BsM (100% or 50% vol/vol solution) was loaded in a 1 mL, 3 mL, or 5 mL syringe on ice, and subsequently transferred to 37 °C for at least 30-60 min to gelate. A predetermined volume of solid matrix was slowly (drop wise) extruded through a 32 G (0.10 mm I.D.) blunt needle and collected into a conical tube containing a predetermined volume of DBPS^+/+^. The mixture was pipetted up and down 10-20 times with a 1 mL pipette tip. The mixture of BsM fragments and DBPS^+/+^ was cooled on ice for 10 minutes before the addition of Col I. The Col I was added to the matrix while on ice, and then the matrix was vortexed for 20-30s. The matrix was centrifuged at 4 °C at 100g for 15-30 seconds to remove bubbles. The fragments were resuspended with a 1 mL pipette tip gently to avoid introducing bubbles. Finally, the matrix was kept on ice until casting or printing. Then, the matrix was incubated for 1-1.5 hours at 37 °C. Although complete list of matrix formulations can be found in **Table S4**, the optimized matrix consisted of BsM fragments generated from 100% (vol/vol) of BsM mixed at a 60:40 ratio with a volume of with Col I and DBPS^+/+^ to form a final, bulk concentration of 60% (vol/vol) BsM (frag.) and 1.5 mg/mL Col I.

### Picrosirus red staining and assessment

Col I fibers were stained following established protocols.^42,43^ Briefly, fixed matrices were stained for 1 hour in a 1 mg/mL picrosirius red solution and washed with two exchanges of 0.5% (vol/vol) solution of acetic acid in de-ionized (DI) water. Images of matrices were taken under brightfield.

### Fragment size analysis

BsM fragments were generated as described except that a 10 mg/mL TRITC-dextran (150 kDa) solution in DBPS^+/+^ was added to the 100% (vol/vol) BsM solution at a ratio of 1:20 before gelation. Fragments were collected in a petri dish of DBPS^+/+^ and immediately imaged by confocal microscopy. Maximum intensity projections were generated, and the fragment areas were segmented and measured in FIJI. Areas >10,000 μm^2^ were observed to be from multiple fragments overlapping and excluded from analysis.

### Rheological measurements

A controlled stress rheometer (DHR-3, TA Instruments, New Castle, DE, USA) with a 25-mm-diameter plate (disposable aluminum, roughened) plate geometry with a gap of 1000 µm was used to measure the rheological properties of the ECM. For time sweeps, a premixed ECM solution was rapidly placed onto the Peltier plate at 4 °C with the temperature subsequently ramped and held at 37 °C in a humidified chamber. The shear storage (G′) and loss (G″) moduli were measured at a frequency of 0.1 Hz and an oscillatory strain (γ) of 0.1%. Rheological testing included frequency sweeps ranging from 10 to 0.01 Hz at 0.1% amplitude also in a humidified 37°C chamber. Shear modulus was reported as the average storage modulus for 3 tests per matrix composition at an oscillation frequency of 0.5 Hz. Shear yield stresses were measured by carrying out amplitude sweeps at 0.1 Hz.

### 2D monolayer assay

ECM substrates for 2D monolayers were prepared in the bottom of a 48-well glass bottom plate. First, glass was coated with 45 µL of 0.1 mg/mL sterile poly-D-Lysine solution in DI water for 1-2 min and aspirated. Wells were washed with sterile MilliQ water, aspirated, and left at room temperature to dry overnight. On the glass region of each well, 45 µL of ECM solution was cast and allowed to crosslink as described above for each type of ECM. To maintain humidity in plates, sterile MilliQ water was added to the outside wells and in between wells. Collagen, and gelbrin matrices were prepared up to 2 days in advance and stored at 4°C in sealed, sterile containers. Dissociated UB cells were seeded on 2D substrates at a density of 385,000 cells / cm^2^ in a 50 µL UBCM-Y droplet and left to adhere for ∼1 hr. Subsequently, 300 µL of UBCM-Y was added to each well. The following day media was replaced with 300 µL of UBCM per well. Samples were fixed on the following day (2 days post seeding) with 10% buffered formalin for 10 min. Fixed samples were permeabilized with 0.1% Triton X and stained with DAPI and ActinGreen 488. Finally, samples were imaged in epifluorescence on a Leica DMIL LED microscope, and analyzed using ImageJ analysis software to segment cells and determine cell area.

### 3D UB budding assay

After aggregation, UPC organoids were embedded in 3D ECM. First, a 200 µL layer of ECM was placed into a 12 mm Transwell culture insert and incubated at 37 °C for ∼1 hr. Then, 200 µL of ECM was mixed with approximately 10-25 organoids in <20 µL of medium and cast on top of the first layer of ECM. The matrices and organoids were incubated for an additional ∼1 hr. Finally, 1.5 mL of UBCM was added to the bottom of the well and changed every 2-3 days. Phase images were collected at day 1 (day of embedding) day 7, day 14, and day 21. At day 21, samples were imaged and fixed. Phase contrast images of organoids were acquired using a 4X or 10X objective (Leica DM IL LED Inverted Laboratory Microscope). Organoid area was quantified by manually tracing the organoid perimeter in FIJI and measuring the organoid area. Fixed samples were stained and imaged to determine marker expression.

### Perfusable 3D ureteric bud tubules and collecting duct models

Custom 3D perfusable chips were fabricated briefly as follows (**Fig. 2B, S4A**). The perfusion platform was fabricated by laser cutting and assembling acrylic sheets and attaching motor apparatus. The metal bases were machined from 0.12” thick stainless steel or aluminum sheets. Gaskets and reservoir lids were acquired from CellTREAT devices. Custom chips were printed from Biomed Clear ink using a FormLabs 3B+ printer. Additional custom components for the chip and platform were printed in Rigid 10k ink using a FormLabs 3B+ printer. Chips were washed in isopropanol for 15 minutes and cured for 1 hour at 60 °C in the Formlabs Form Cure. Chip components were either sterile packed and autoclaved or sterilized by 70% ethanol exposure. A bill of materials, CAD files, and detailed fabrication protocols will be provided upon request.

Sterilized components of the perfusion assembly were assembled in a biosafety cabinet (**Fig. S4A**). Before tubing was added, a Ø1 mm stainless steel pin was inserted in the outlet and completely immersed with ECM solution in the inner reservoir (**Fig. 2B**). The matrix was incubated at 37°C for 1hr to gelate. Afterwards, UBCM-Y with 1x antibiotic-antimycotic was added to each basal chamber to hydrate and equilibrate with the matrix at humidified 37 °C for at least 2 hours. Then, the 1 mL of UBCM-Y was transferred to the apical chamber and the pin was removed through the outlet, leaving a bubble-free, medium-filled channel embedded within the gel. Microbore silicone tubing (0.50-mm inner diameter) connected to PharMed BPT tubing (0.25 mm ID) via custom adapters to the apical or basal chambers. The large gear and rollers were attached to clamp the tubing shut when not in use. Channels were seeded with UB cells (day 21-35), which were dissociated as described above and concentrated in UBCM-Y to a density of ∼100 x 10^6^ cells/mL. Medium was removed from both the apical and basal reservoirs (but not the channel), and 25 µL of cell suspension was injected into the channel via the reservoir outlet. The chip was left upright for 10 minutes, then rotated from side to side for 10 minutes each to coat the channel. Tubing was reattached, medium was added to the apical and basal reservoirs, and cells were left to adhere at 37°C and 5% CO2 overnight. The next day, nonadherent cells were flushed out with UBCM-Y and the chip reservoir medium was refreshed.

Our platform is highly modular and can be easily transferred from incubators into biosafety cabinets for sterile medium changes without requiring tubing to be unclipped from the peristaltic pump and adjusted for transport. In our experience, tubing adjustments during transfer are time consuming and often detrimental to soft matrices; pressure changes in the lines can cause channels to rapidly burst or collapse (data not shown).

Cells were left to form confluent monolayers (7 – 14 days) in UBCM-Y, after which the chips were connected to the pump platform (except static chip controls). The luminal flow rate of the chip system could be adjusted within a range of 2.5-66 μL/min by dialing the rotations per minute (RPM) of the pump motor (**Fig. S4B**). Medium was changed to either UBCM or CD at the start of perfusion. Constructs were continuously perfused using a unidirectional peristaltic flow rate of 5 µL/min (0.006 dyn/cm^2^). After 1 day of perfusion at 5 µL/min, flow rate was either maintained at 5 µL/min (low flow) or increased to 66 µL/min (0.08 dyn/cm^2^; high flow). In line with native CD cells which are sensitive to shear stresses in the range of ∼0.01 dyn/cm^2^ to 0.1 dyn/cm^2^ ^44–46^. Chip reservoir and perfusion reservoir mediums were changed every 2-3 days and cultured from 2 days to 2 weeks. All media on chip included amphotericin B at 0.25 ug/mL. Medium volumes in the apical and basal reservoirs were maintained at 600 µL and 900 µL, respectively, for regular maintenance. Tubules on chip were imaged in phase using a Leica DMIL LED microscope equipped with 4x and 10x objectives. Diameters and bud lengths were measured manually in FIJI.

We reproducibly fabricated and perfused tubules on chip in UB or CD medium in N=3 independent experiments with 33 of 36 total tubules successfully cultured to the experimental endpoint (**Fig. S8**). CD differentiation of UB organoids and UB tubules on chips was performed as per the protocol of Zeng and Huang *et al.*^9^. Briefly, UBCM medium replaced by CD differentiation medium (**Table S4**) was added. CD differentiation was initiated on oUB-lined channels and whole organoid controls 7 – 14 days post seeding oUBs on chip. To ensure thorough washout of UBCM on chip, CD medium was refreshed twice on the first day of perfusion and again the next day. Medium was refreshed every 2-3 days for 7 days.

### Embedded bioprinting

Filamentary and branching UPC networks were printed using a custom-designed, multimaterial 3D bioprinter equipped with four independently addressable printheads mounted on a three-axis, motion-controlled gantry (AGB 10000, Aerotech Inc., Pittsburgh, PA, USA) equipped with a custom-built syringe pump mounted on the 3D printer. The syringe pump was operated by an Arduino microcontroller and a stepper-motor driver. All later prints in transwells, custom chambers, and chips were performed in a custom low-cost open source printer mounted with a custom chilled sample holder within a biosafety cabinet.^39^ UB cell slurries were loaded into a 100 µL glass syringe. A metal nozzle, with a length of 1.5 inches and an inner diameter of 0.25 mm, was fitted to the bottom of the syringe. The glass syringe was then loaded onto the syringe pump that was mounted onto the 3D printer, and printer motion was controlled with custom G-code. An extrusion volumetric flow rate of 0.80 μL/s and nozzle translation speed of 60 mm/min were selected for printing.

### Channel stretch test

To determine the channel stretch sensitivity to reservoir medium volume, non-epithelialized channels were cast, and the apical and basal medium volumes were varied without perfusion. The apical volumes were varied between 0 and 1.2 mL in 0.4 mL increments and the basal volumes were varied between 0 and 3 mL in 1 mL increments. These volumes were predetermined to vary the height of the reservoir volumes by ∼5 mm increments in both apical and basal chambers. For each measurement, the appropriate volume of medium was added to each reservoir and equilibrated for 5 minutes to allow stretching or shrinking to occur. At 5 minutes, channel was imaged using a Leica DMIL LED microscope equipped with 4x objective in phase before medium was removed. Channel diameters were measured in FIJI, and data represent single channel measurements from 4 independent experiments.

### Barrier assay

A stock FITC-inulin solution at 1 mg/mL in DBPS^+/+^ was prepared and diluted to 100 µg/mL in UBCM. Chips with and without confluent epithelialized channels were perfused with FITC-inulin containing medium for 60 minutes at a volumetric flow rate of 33 µL/min. Channels were imaged in phase and epifluorescence using a Leica DMIL LED microscope equipped with a 4x objective before the addition of FITC inulin, after 30 minutes of perfusion, and after 60 minutes of perfusion. Fluorescence intensity profiles were analyzed in ImageJ. Intensity profiles were normalized to the average intensity of the profile before addition of the FITC-inulin by subtracting the average value from each data point along the profile. Data represent N=3 independent experiments.

### RNA isolation and Nanostring analysis

For each condition, ∼10 organoids or 1-2 channels were collected. To minimize matrix, excess matrix was sliced off the channel on top, bottom, and both sides by scalpel. Samples were transferred to epitubes and rinsed with DBPS^+/+^. RNA was isolated using the RNeasy Plus Mini Kit according to manufacturer’s instructions. RNA concentration was assessed using both the Qubit and Nanodrop 1000 spectrophotometer. Probes of a custom kidney gene panel for NanoString analysis was utilized, using either the full panel of 172 key genes or an abbreviated panel of the first 48 key genes (**Table S5**). RNA was quantified with a NanoString nCounter Elements™ with according reagents as per the manufacturer’s instructions, using 50-100 ng RNA per run. RNA was hybridized with probe pools, hybridization buffer, and TagSet reagents in a total volume of 30 μL and incubated at 67 °C for 20 h. Quantification data was analyzed by nSolver software using a custom advanced analysis. No low count data was omitted in initial analysis. *ACTB*, *GAPDH*, and *TUBB* were marked as housekeeping genes. After processing the data in nSolver, low count data was removed from further analysis. Low count genes were defined as genes in which the mRNA count was below threshold for at least one replicate in all experimental conditions. Data was plotted with R.

### Immunostaining and imaging

Samples were briefly washed with DPBS^+/+^and subsequently fixated in 10% buffered formalin solution for 1h at RT. The fixative was aspirated, and samples were washed 3x in DPBS^+/+^. Chip samples were kept whole or sectioned with a scalpel. Subsequently, samples were blocked and permeabilized in DPBS^+/+^ supplemented with 0.125% v/v Triton X-100 and 1% v/v donkey serum for 24 h. Samples were exposed to primary antibodies (**Table S2**) for 24 h at 4 °C in a staining solution (0.5% w/v BSA and 0.125% v/v Triton X-100 in DPBS^+/+^). Post incubation, samples were washed with DPBS^+/+^. A cocktail of secondary antibodies and stains (i.e. DAPI, phalloidin) (**Table S2**) was incubated with the samples for 24 h at 4 °C in staining solution. Finally, samples were washed 3x in DPBS^+/^. Images were acquired with 5x, 10x, 20x, and 40x water dipping objectives on a Zeiss LSM 710 confocal microscope. Images, 3D projections, stacks, and rendered views were generated in FIJI and Imaris. In some cases specified in figure legends, samples were treated with Easy Index before imaging with air objectives. ImageJ was used for processing and quantification.

### Statistical analysis

Prior to experimentation, the required number of biological replicates to achieve power of 0.8 were estimated using G*Power using previous or preliminary data. Statistical significance was determined via indicated tests using GraphPad/Prism or Nsolver. Detailed statistical methods are listed in figure legends including: exact value of N and what N represents, statistical test used for comparison, definition of center, dispersion and precision measures, and definition of significance.

## Acknowledgements

This work was supported by funding from the Wellcome LEAP Human Organ Physiological Engineering (HOPE) program, the NIH Re(Building) a Kidney Consortium (NIH UC2DK126023), the NIH F32 Ruth L. Kirschstein National Research Service Award for Individual Postdoctoral Fellows (KJW, DK131821), and Dutch Research Council Rubicon grant (RG, 019.201EN.005). The authors would like to thank staff at the Bauer Core at Harvard University for assistance with cell sorting, personnel at the machine shops of Weiss institute and School of Engineering and Applied Sciences at Harvard University for part fabrication of the modular chip with integrated pump, Collin Fullen for early contributions on chip validation, Alana Mermin-Bunnell for aiding in 3D printer assembly, Paul Stankey and Ben Fichtenkort for aiding in the generation of the fluorescent iPSC line, and Dr. Alexander Ainscough for assistance with Nanostring Analysis. Finally, we thank Alice Chen and Ben Shepard for useful discussions.

