## Supplemental Figures and Tables for "Perfusable 3D models of ureteric bud and collecting duct tubules"

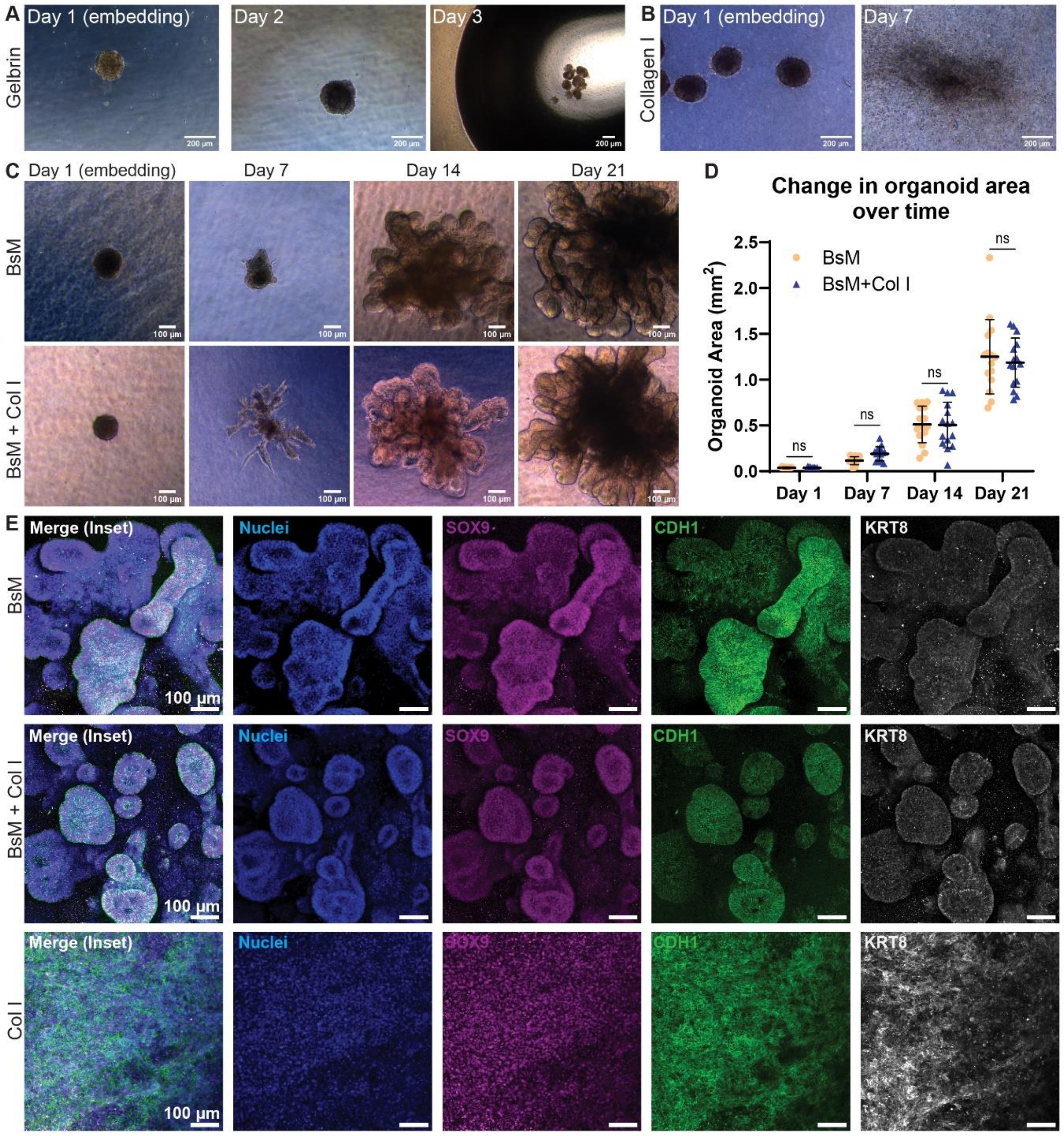


**Supplementary Figure 1. Uretic bud organoids cultured in different matrices**. UPCs 1 day post-aggregation were embedded in matrices differentiated for up to 21 days into UB organoids. Phase images of aggregates in **A**) Gelbrin, scale bar = 200 µm **B**) Collagen I, scale bar = 200 µm, **C**) BsM (top), BsM - Col I (bottom), scale bar = 100 µm. **D**) Organoid area quantification of bright field images in BsM (orange) and BsM - Col I (blue) over time. Non-significant (ns) from statistical analysis by two-way ANOVA followed by Sidak's multiple comparisons test. Organoids within matrices were compared at each timepoint. Data points represent areas of 15 individual N=15 organoids from 3 independent experiments. Error bars denote mean and standard deviation. **E**) Immunofluorescent images of UB organoids in BsM, BsM - Col I and Col I matrices (from top to bottom). Stained for nucleus (blue), SOX9 (magenta), CDH1 (green), KRT8 (white) at 21 days of differentiation. Region of interest from Figure 1D. Scale bar = 100 µm.

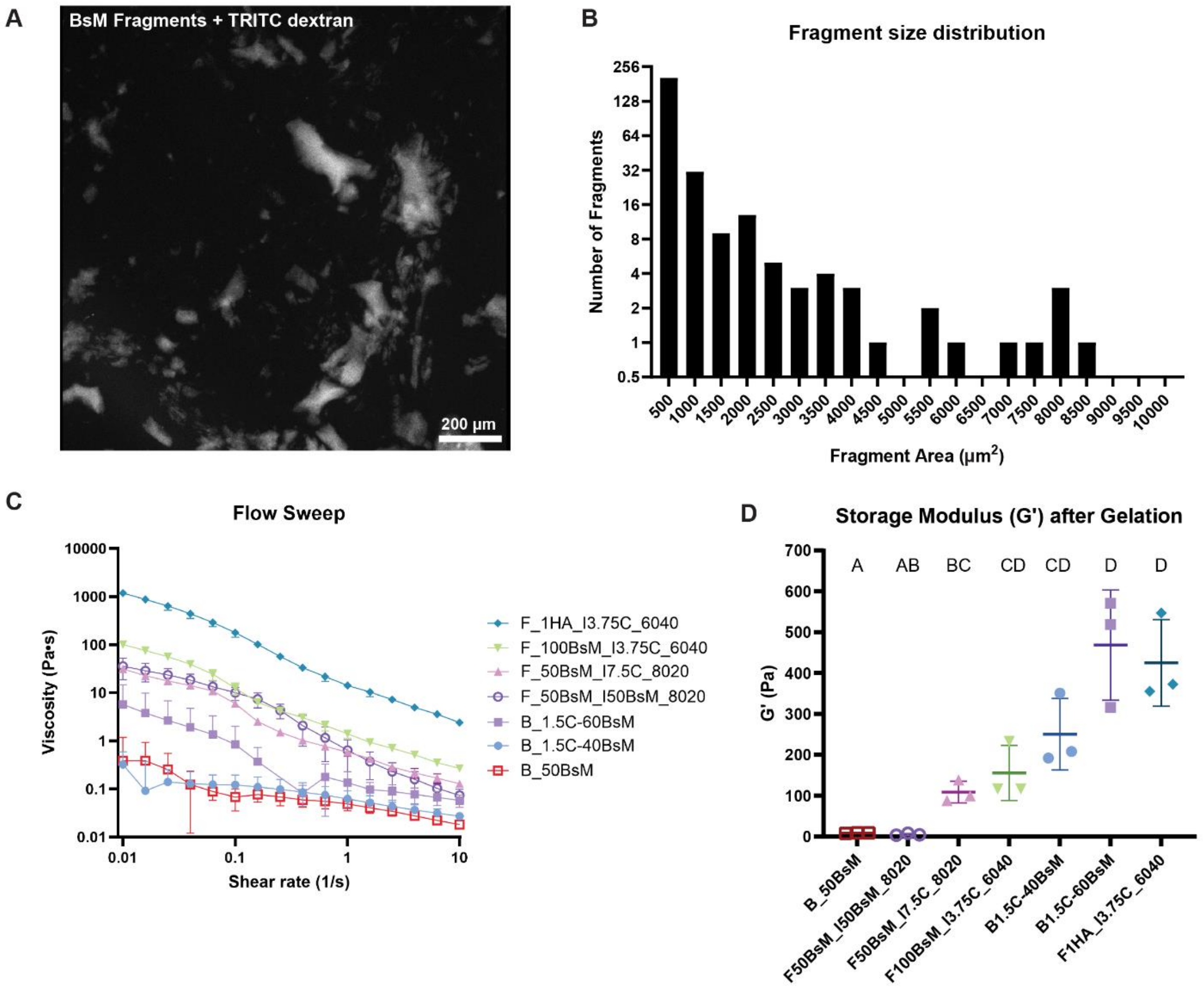


**Supplementary Figure 2. Fragmented matrix characterization.** **A**) Max intensity projection fluorescent image of BsM fragments loaded with TRITC-dextran. Scale bar = 200 µm. **B**) Fragment size distribution of BsM fragments. Histogram counts are the sum from N=3 independent fragmentation batches. **C**) Flow sweeps at 4 °C before collagen crosslinking and **D**) storage modulus after collagen crosslinking at 37 °C. Error bars denote mean and standard deviation. Letters represent statistical families defined as $p<0.05$ by one-way ANOVA followed by Tukey’s test for multiple comparisons. Data points represent N=3 independent experiments defined as independent matrix batches. Matrix compositions denoted in legend are defined in Table S4.

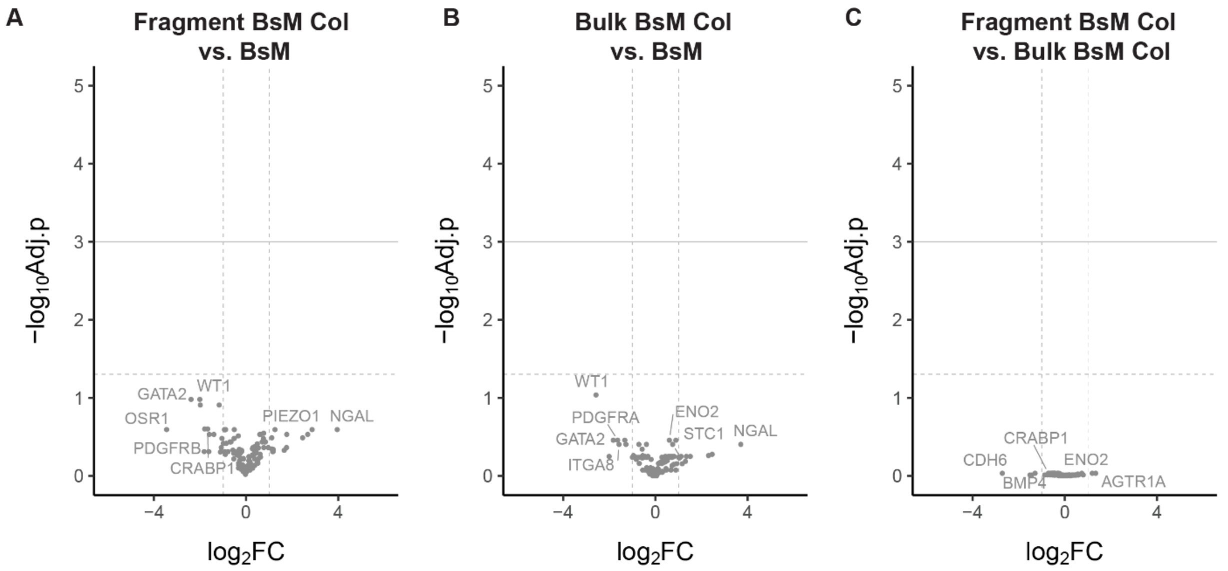


**Supplementary Figure 3. RNA expression of UB organoid in matrices**. **A**) BsM (frag), **B**) BsM-Col I, **C**) BsM (frag.) - Col I vs BsM. Color corresponds to non-significantly expressed genes (gray), dashed horizontal line = adjusted $p<0.05$, solid horizontal line = adjusted $p<0.01$, vertical line = 2-fold change. Datapoints represent individual genes from N=3 independent experiments where p-values are adjusted by the Benjamini-Hochberg correction.

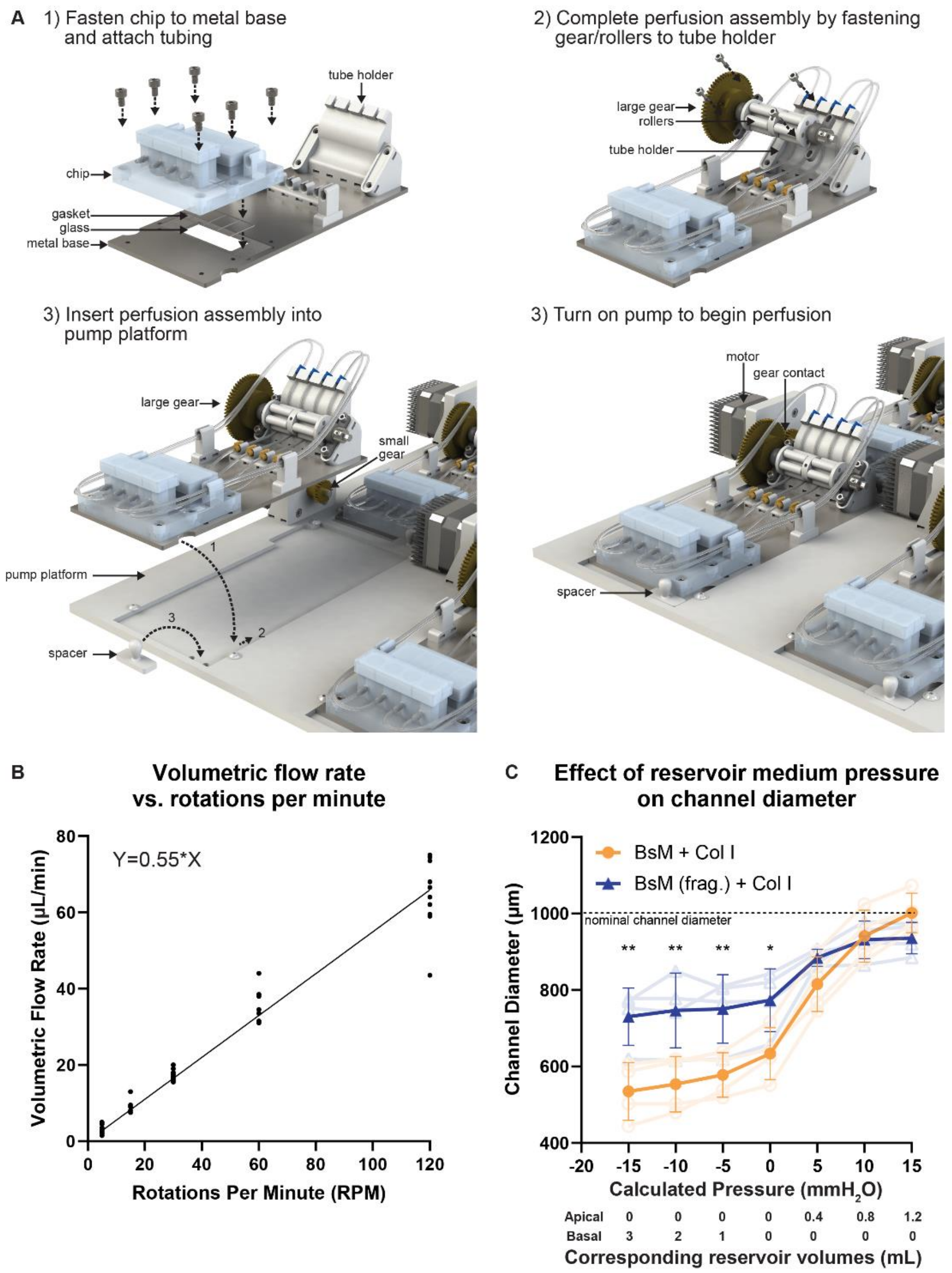


**Supplementary Figure 4. Assembly and characterization of modular chip with integrated pumps. A**) Assembly schematic of modular chip. **B**) Correlation of measured volumetric flow rate relative to motor rotations per minute and the resulting formula thereof. Datapoints represent measurements from 10 individual channels collected over 4 independent experiments. Line and equation generated by simple linear regression analysis. **C**) Channel diameter in response to pressure differences created by different volumes of medium in the apical and basal reservoirs for each matrix. BsM-Col I (orange= average, light orange= individual replicates) and BsM (frag.) - Col I (blue= average , light blue= individual replicates). Error bars denote mean and standard deviation. N=4, *p<0.05, **p<0.01 by two-way ANOVA followed by Sidak's multiple comparisons test.

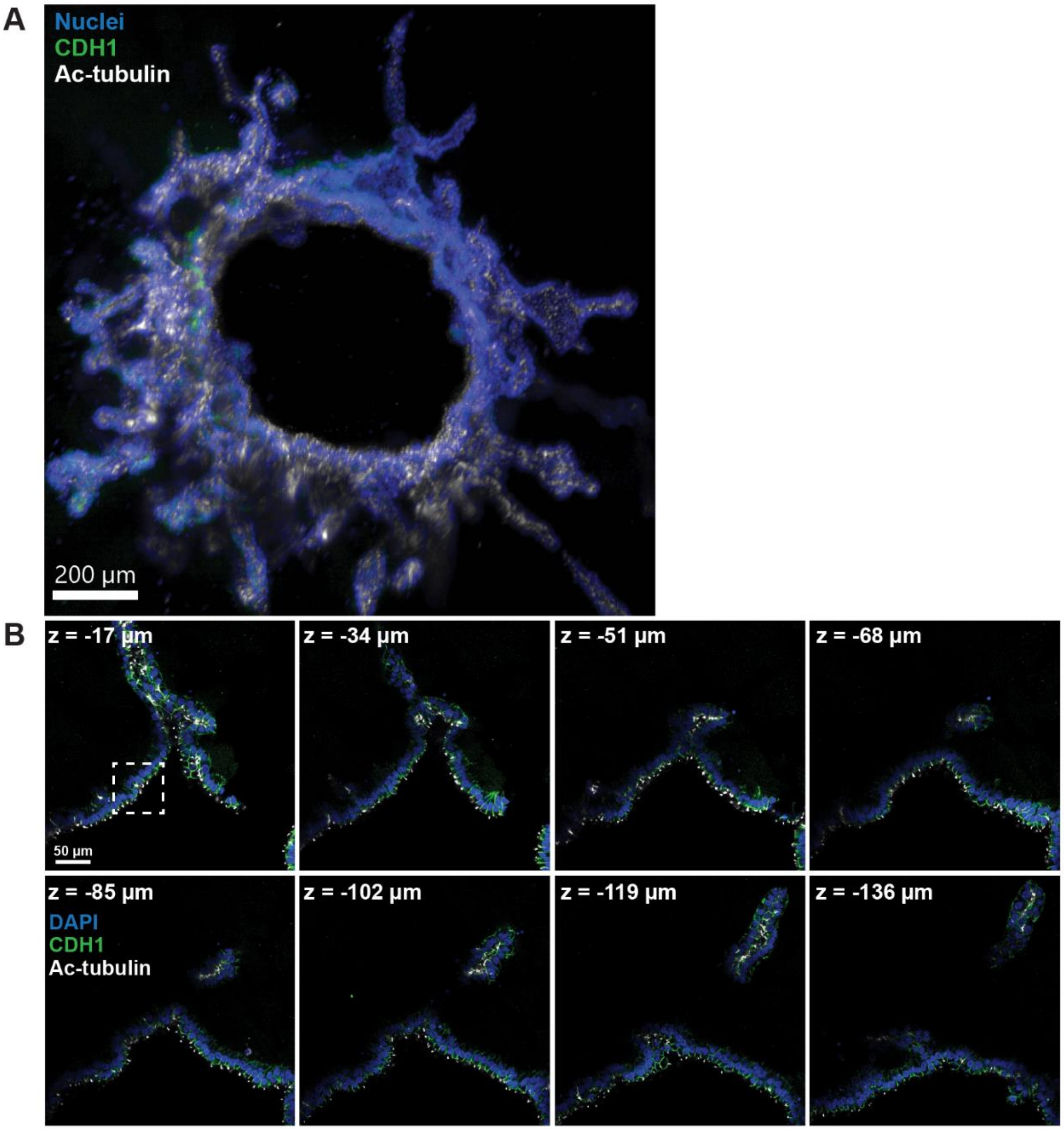


**Supplementary Figure 5. 3D UB tubule under high flow. A)** Z-projection of UB tubule imaged with easy-index. Cells stained for nuclei (blue), CDH1 (green) Ac-tubulin (white). Scale bar = 200 µm. Sample corresponding to image Figure 2D-E. **B**) Montage of single z-slices of same region of interest at different depths. Scale bar = 50 µm.

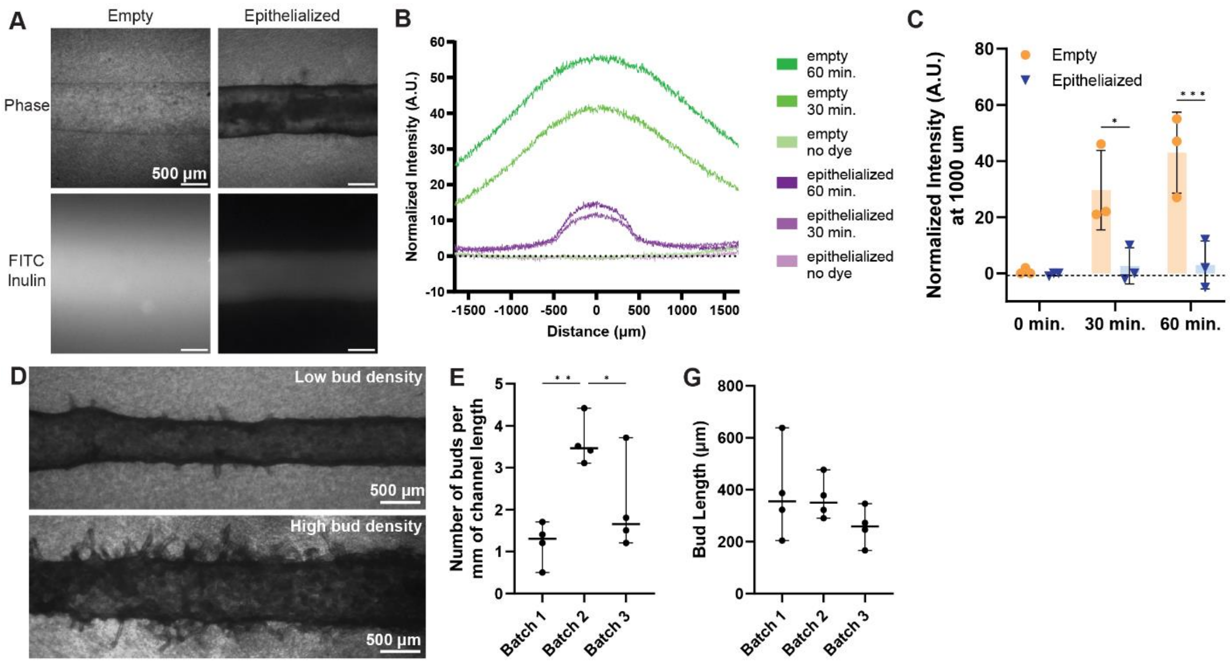


**Supplementary Figure 6. Characterization of perfusable 3D UB tubules.** **A**) Phase images of empty and epithelialized (oUBs) channel (top) with corresponding fluorescent images of FITC-inulin (bottom) after 60 minutes of perfusion at a volumetric flow rate of 33 µL/min. Scale bar = 500 µm. **B**) FITC-inulin fluorescence distribution over the image at 0, 30, and 60 min for both conditions (green = empty channel; purple = epithelialized channel). Traces represent an average over N=3 independent experiments. **C)** Fluorescence at 1000 µm from the channel at 0, 30, and 60 min (orange = empty channel; blue = epithelialized channel). Datapoints represent measurements from individual channels from 3 independent experiments. *****p<0.05, ***p<0.001 by two-way ANOVA followed by Sidak's multiple comparisons test in which empty and epithelialized channels at each time point were compared. **D**) Phase images depicting batch-to-batch variability found in UB budding morphology on chip. Scale bar = 500 µm. Quantification of **E**) bud numbers along the length of the channel and **F**) maximum bud length from channel. Datapoints represent counts from individual channels from 4 independent experiments. *****p<0.05, **p<0.01 by one-way ANOVA followed by Tukey's multiple comparisons test. Error bars denote median and range.

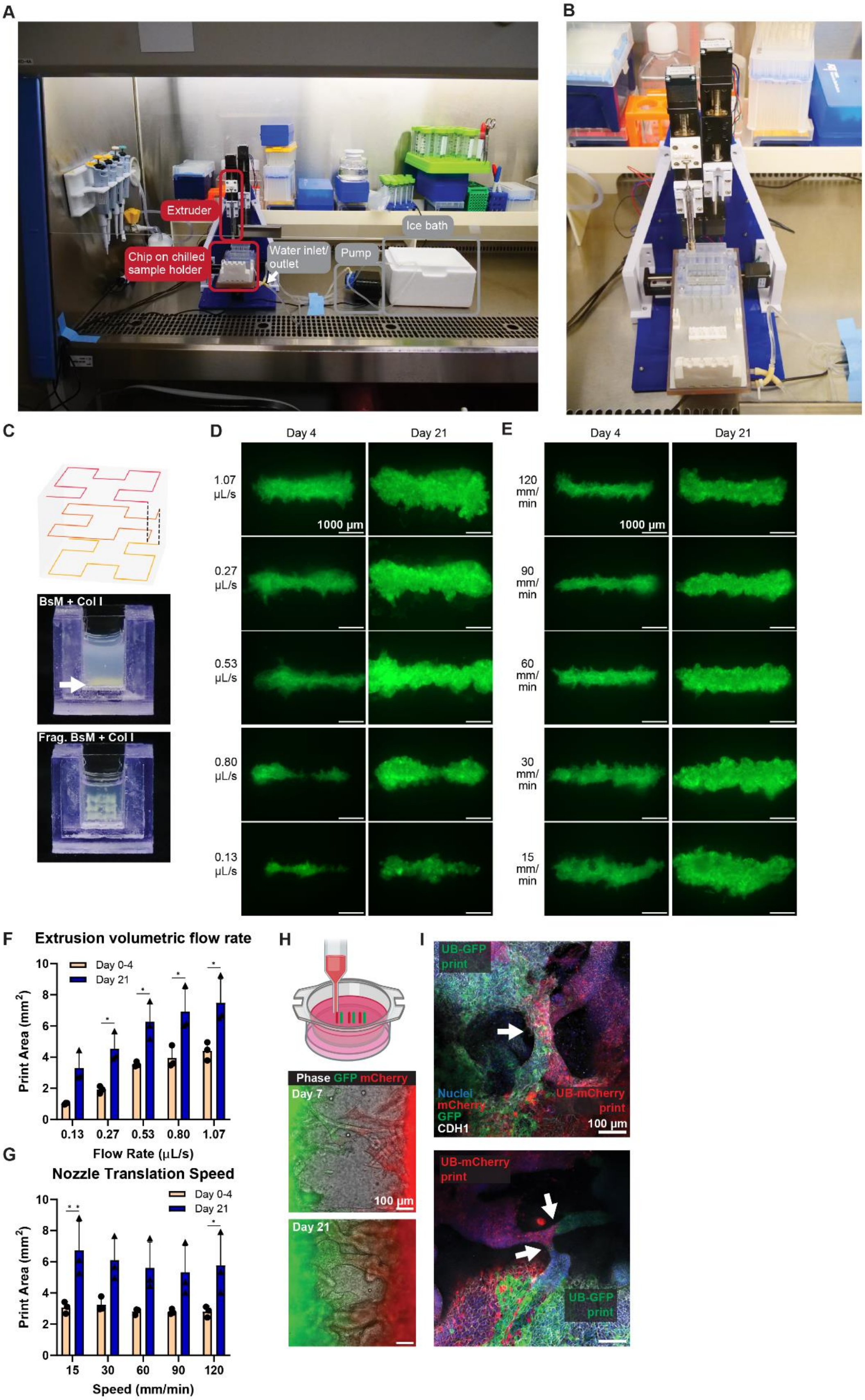
A
Extruder
Chip on chilled sample holder
Water inlet/ outlet
Pump
Ice bath
B
C
BsM + Col I
Frag. BsM + Col I
D
Day 4
Day 21
1.07 µL/s
0.27 µL/s
0.53 µL/s
0.80 µL/s
0.13 µL/s
1000 µm
E
120 mm/ min
90 mm/ min
60 mm/ min
30 mm/ min
15 mm/ min
F
Extrusion volumetric flow rate
Day 0-4
Day 21
Print Area (mm2)
Flow Rate (µL/s)
0.13
0.27
0.53
0.80
1.07
G
Nozzle Translation Speed
Speed (mm/min)
15
30
60
90
120
H
Phase GFP mCherry
Day 7
Day 21
100 µm
I
UB-GFP print
UB-mCherry print
Nuclei
mCherry
GFP
CDH1
100 µm

**Supplementary Figure 7. Sterile embedded bioprinting set-up and validation.** **A**) Picture of in-hood printing set-up[1] with custom holder for on chip printing. **B)** Configuration for printing directly into modular perfusion platform. **C**) Validation of BsM (frag.) - Col I for bioprinting. Print path (top), embedded printing in a BsM - Col I matrix where arrow indicates extruded cells that have settled to the bottom of print chamber (middle), and embedded printing in BsM (frag.) - Col I in which printed cells are supported by matrix (bottom). **D)** Epifluorescence of extruded GFP-expressing UB cells relative to extrusion volumetric flow rate. Scale bar = 1000 µm. **E)** Epifluorescence of extruded GFP-expressing UB cells relative to nozzle speed. Scale bar = 1000 µm. Quantification of cell area as function of **F)** extrusion volumetric flow rate and **G)** nozzle translation speed at days 4 and 21 post print**.** *$p<0.05$, **$p<0.01$ by two-way ANOVA followed by Sidak's multiple comparisons test where print area was compared between time points within each print condition. Error bars depict mean and standard deviation. Data points represent measurements of single print lines from N=3 independent experiments. **H**) Schematic overview of printing 4 mm alternating GFP- or mCherry-expressing UPCs in transwells (top), phase and epifluorescence images of tubules fusing over time (bottom). Phase (gray), GFP (green), mCherry (red). Scale bar = 100 µm. **I**) Immunofluorescence images of UB tubules 21 days post print interconnecting between adjacent lines showing nuclei (blue), mCherry (red), GFP (green), and CDH1 (white). GFP and mCherry cells exhibited mixing at some fusion sites (top) but not others (bottom). Scale bar = 100 µm.

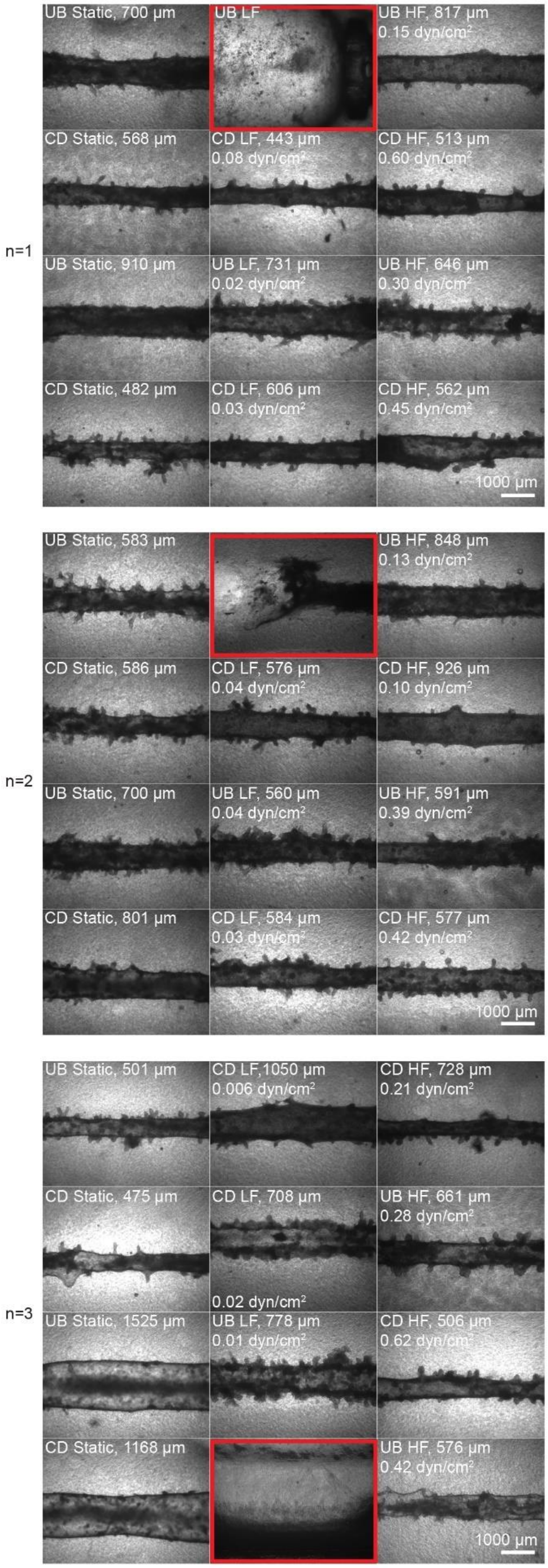

UB Static, 700 µm
UB LF
UB HF, 817 µm
0.15 dyn/cm²
CD Static, 568 µm
CD LF, 443 µm
0.08 dyn/cm²
CD HF, 513 µm
0.60 dyn/cm²
n=1
UB Static, 910 µm
UB LF, 731 µm
0.02 dyn/cm²
UB HF, 646 µm
0.30 dyn/cm²
CD Static, 482 µm
CD LF, 606 µm
0.03 dyn/cm²
CD HF, 562 µm
0.45 dyn/cm²
1000 µm
UB Static, 583 µm
UB HF, 848 µm
0.13 dyn/cm²
CD Static, 586 µm
CD LF, 576 µm
0.04 dyn/cm²
CD HF, 926 µm
0.10 dyn/cm²
n=2
UB Static, 700 µm
UB LF, 560 µm
0.04 dyn/cm²
UB HF, 591 µm
0.39 dyn/cm²
CD Static, 801 µm
CD LF, 584 µm
0.03 dyn/cm²
CD HF, 577 µm
0.42 dyn/cm²
1000 µm
UB Static, 501 µm
CD LF,1050 µm
0.006 dyn/cm²
CD HF, 728 µm
0.21 dyn/cm²
CD Static, 475 µm
CD LF, 708 µm
0.02 dyn/cm²
UB HF, 661 µm
0.28 dyn/cm²
n=3
UB Static, 1525 µm
UB LF, 778 µm
0.01 dyn/cm²
CD HF, 506 µm
0.62 dyn/cm²
CD Static, 1168 µm
UB HF, 576 µm
0.42 dyn/cm²
1000 µm

**Supplementary Figure 8. 3D UB-to-CD tubule differentiation**. Brightfield images of tubules produced under different experimental conditions, channel diameter at most narrow point and calculated fluid shear stress experienced at the channel wall. Cells were cultured for 21 days after seeding on chip. Red-lined images highlight failed channels. Tubule failure modes included unexplained tubule swelling, tubule collapse, and technical error leading to leakage.  Scale bar = 1000 µm.

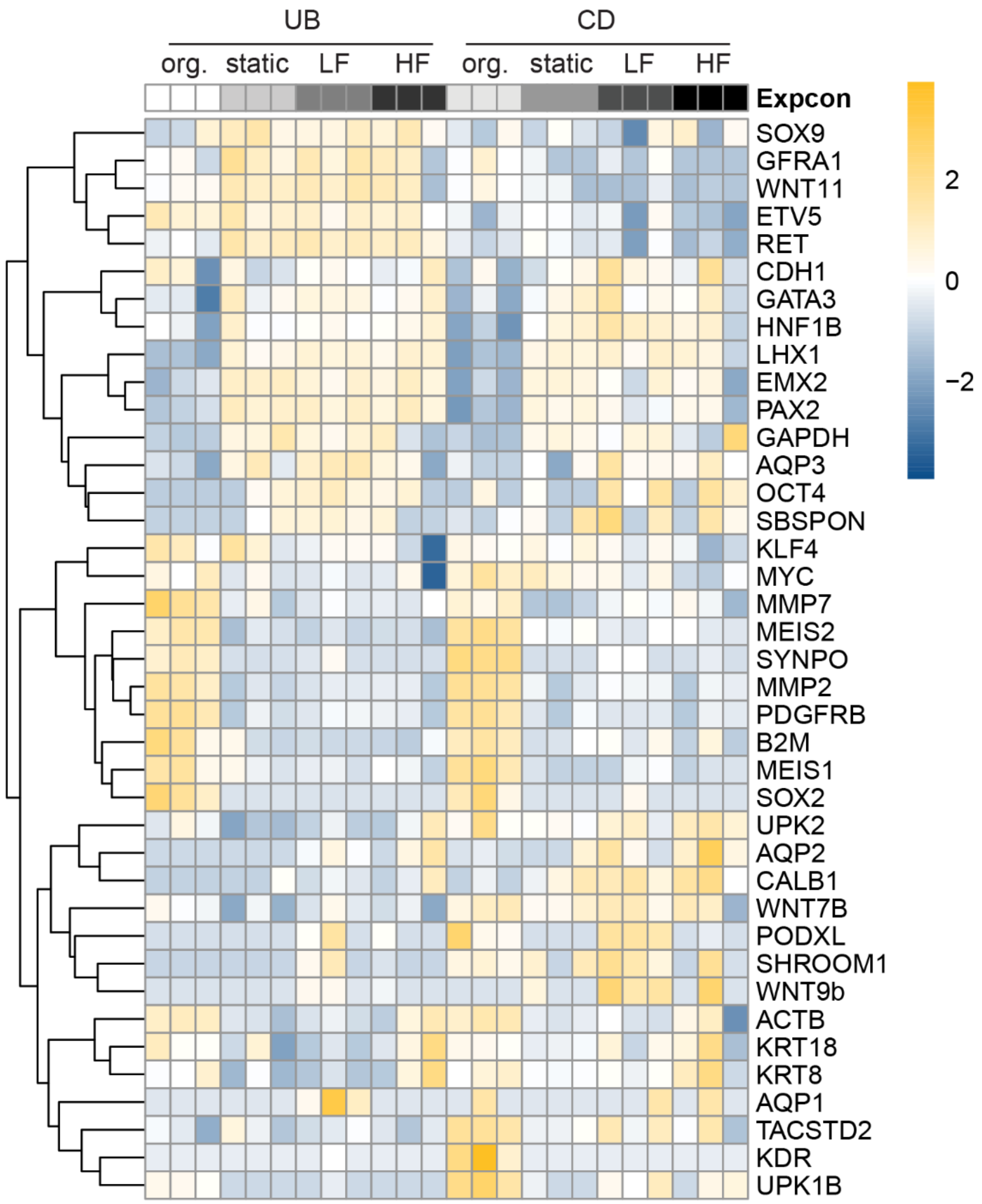


**Supplementary Figure 9. Heatmap of RNA expression.** Heatmap of RNA expression of all investigated genes with counts that passed criteria for analysis. Colors correspond to up- (orange) and down- (blue) regulated genes scaled to the standardized z-score. Conditions from left to right: UB embedded organoid (org.), static UB-on-chip, low flow UB on-chip, high flow UB on-chip, CD embedded organoid (org.), static CD on-chip, low flow CD on chip and high flow CD on chip. All samples were cultured for 21 days after seeding on chip. Columns within each condition represent N=3 independent experiments.

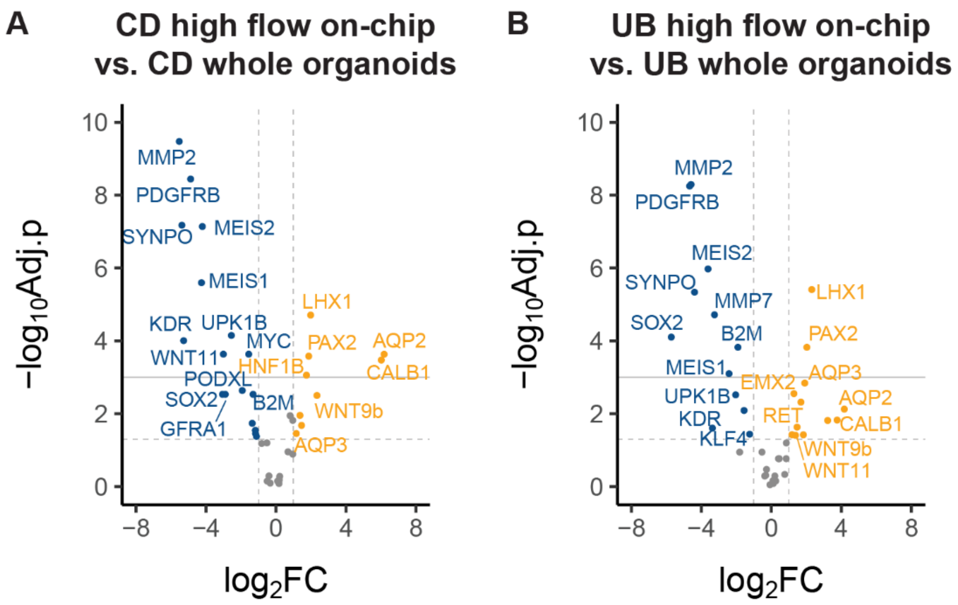


**Supplementary Figure 10. Differential RNA expression HF vs. whole organoids.** Differential RNA expression of all investigated genes with counts that passed criteria for analysis in tubules on chip compared to embedded whole organoids depicted in volcano plots. **A**) High flow CD on chip relative to embedded CD organoids. **B**) High flow UB on chip relative to embedded UB organoids. Colors correspond to non-significantly expressed genes (gray), upregulated genes (orange), and downregulated genes (blue). Dashed horizontal line = adjusted $p<0.05$, solid horizontal line = adjusted $p<0.01$, vertical line = 2-fold change. Data points represent individual genes from N=3 independent experiments where p-values are adjusted by the Benjamini-Hochberg correction.

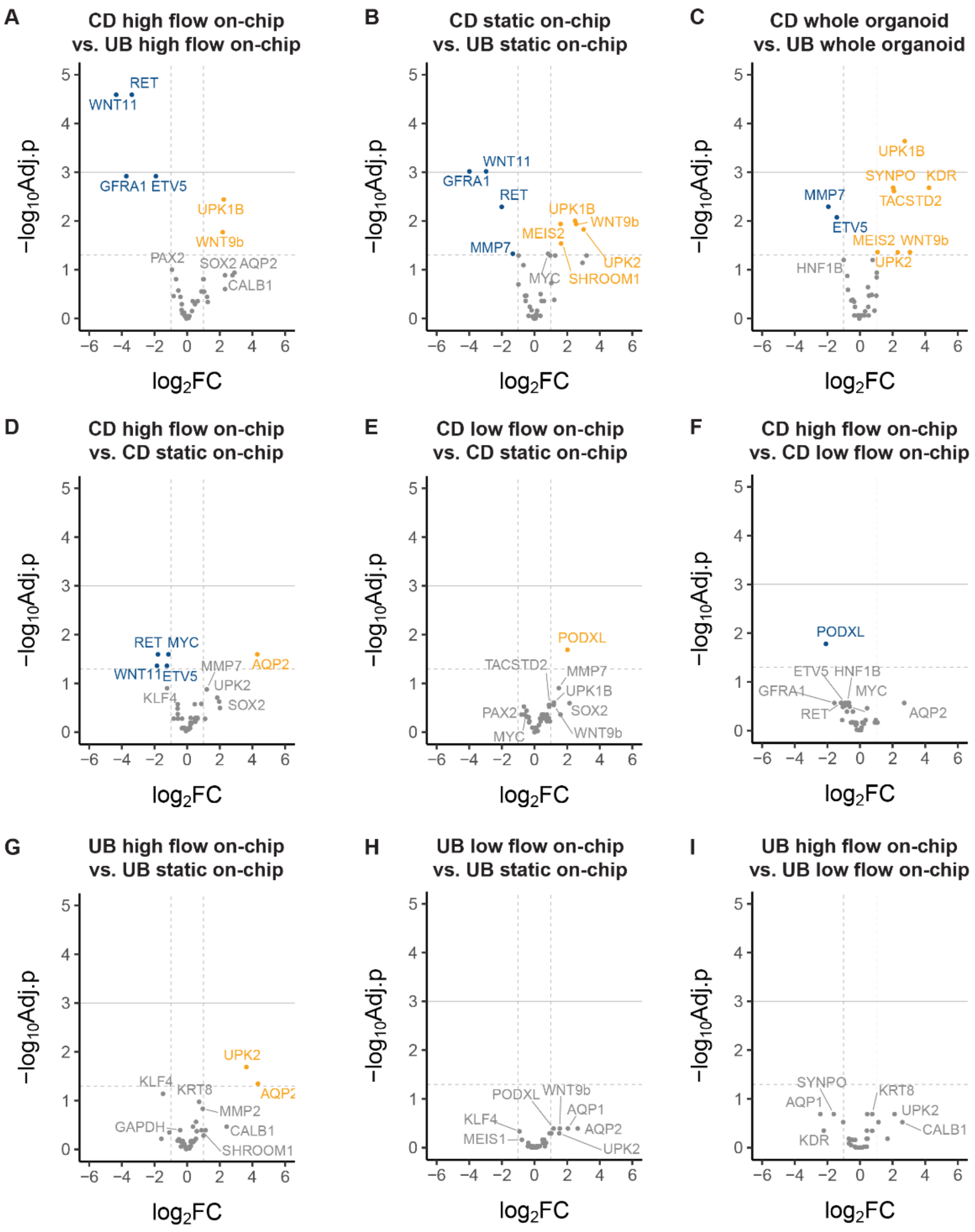


**Supplementary Figure 11. Gene expression across multiple condition comparisons.** Differential RNA expression of all investigated genes with counts that passed criteria for analysis depicted in volcano plots. Throughout figure: Colors correspond to non-significantly expressed genes (gray), upregulated genes (orange), and downregulated genes (blue). Dashed horizontal line = adjusted $p<0.05$, solid horizontal line = adjusted $p<0.01$, vertical line = 2-fold change. Datapoints represent individual genes from N=3 independent experiments where p-values are adjusted by the Benjamini-Hochberg correction. **A)** CD high flow on chip relative to UB high flow on chip, **B)** CD static on chip relative to UB static on chip, **C**) CD embedded whole organoids relative to UB embedded whole organoids, **D)** CD high flow on chip relative to CD static on chip, **E)** CD low flow on chip relative to CD static on chip, **F)** CD high flow on chip relative to CD low flow on chip, **G)** UB high flow on chip relative to UB static on chip, **H)** UB low flow on chip relative to UB static on chip, and **I)** UB high flow on chip relative to UB low flow on chip.

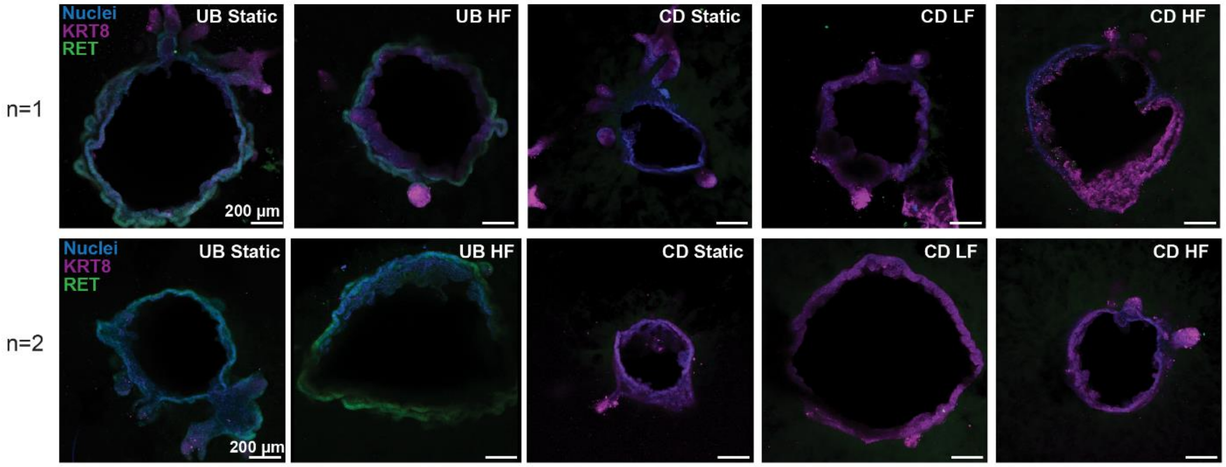


**Supplementary Figure 12. Maximum intensity projection immunofluorescence cross-sectional images of UB and CD on chips**. From left to right, static UB on chip, high flow UB on chip, static CD on chip, low flow CD on chip, and high flow CD on chip. Nuclei (blue), KRT8 (magenta), RET (green). Scale bar = 200 µm. Cells cultured for 14 days after seeding on chip followed by 7 additional days with or without CD medium. N=2 independent experiments. HF = high flow, LF = low flow.

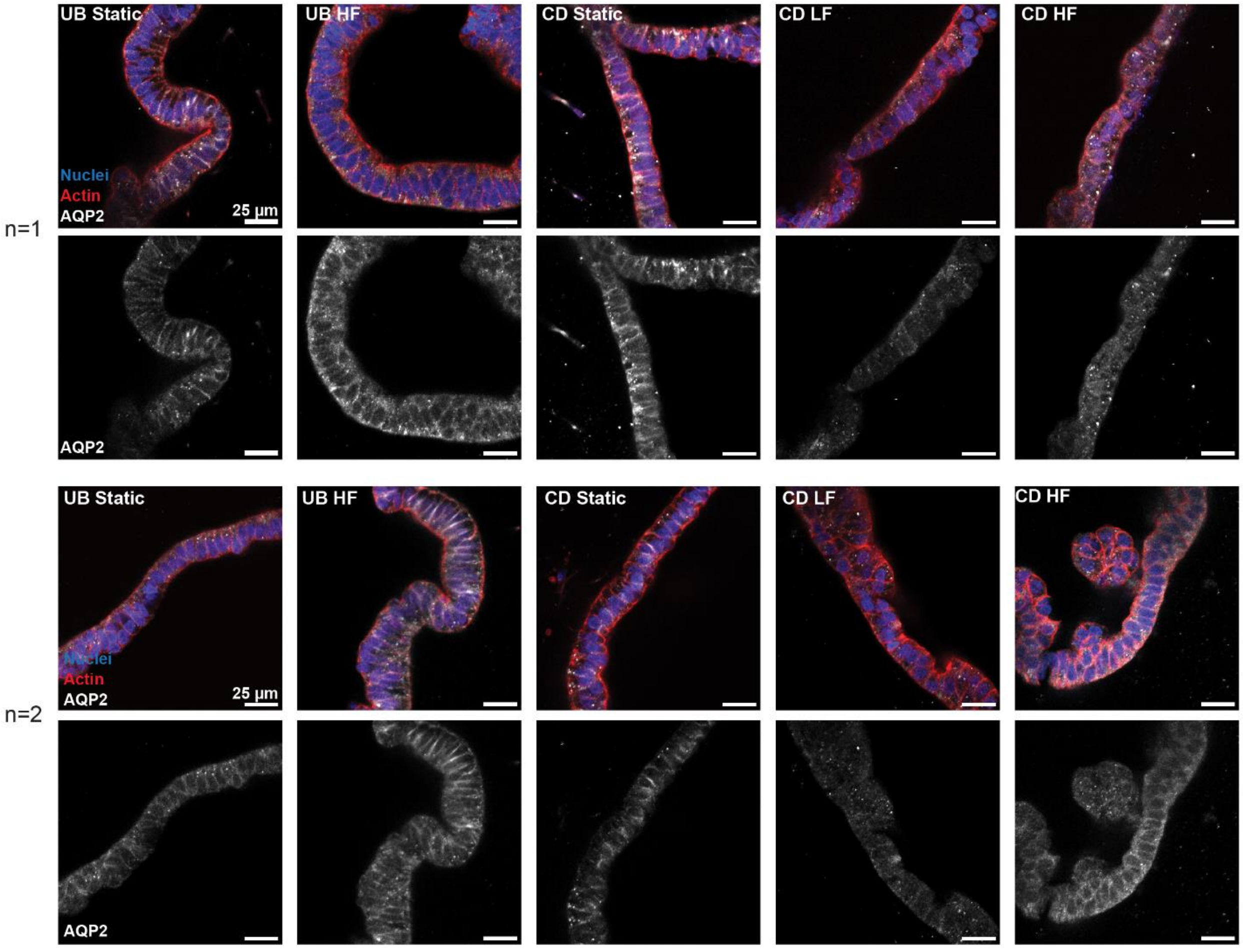


**Supplementary Figure 13. Cross-sectional immunofluorescence images of 3D UB and CD tubules (single z-slices)**. From left to right, static UB on chip (UB static), high flow (HF) UB on chip, static CD on chip, low flow (LF), and HF CD-on chip. Top row: nuclei (blue), actin (red), and AQP2 (white). Bottom row: AQP2 staining only. Scale bar = 25 µm. Cells cultured for 14 days after seeding on chip followed by 7 additional days with or without CD medium. N=2 independent experiments.

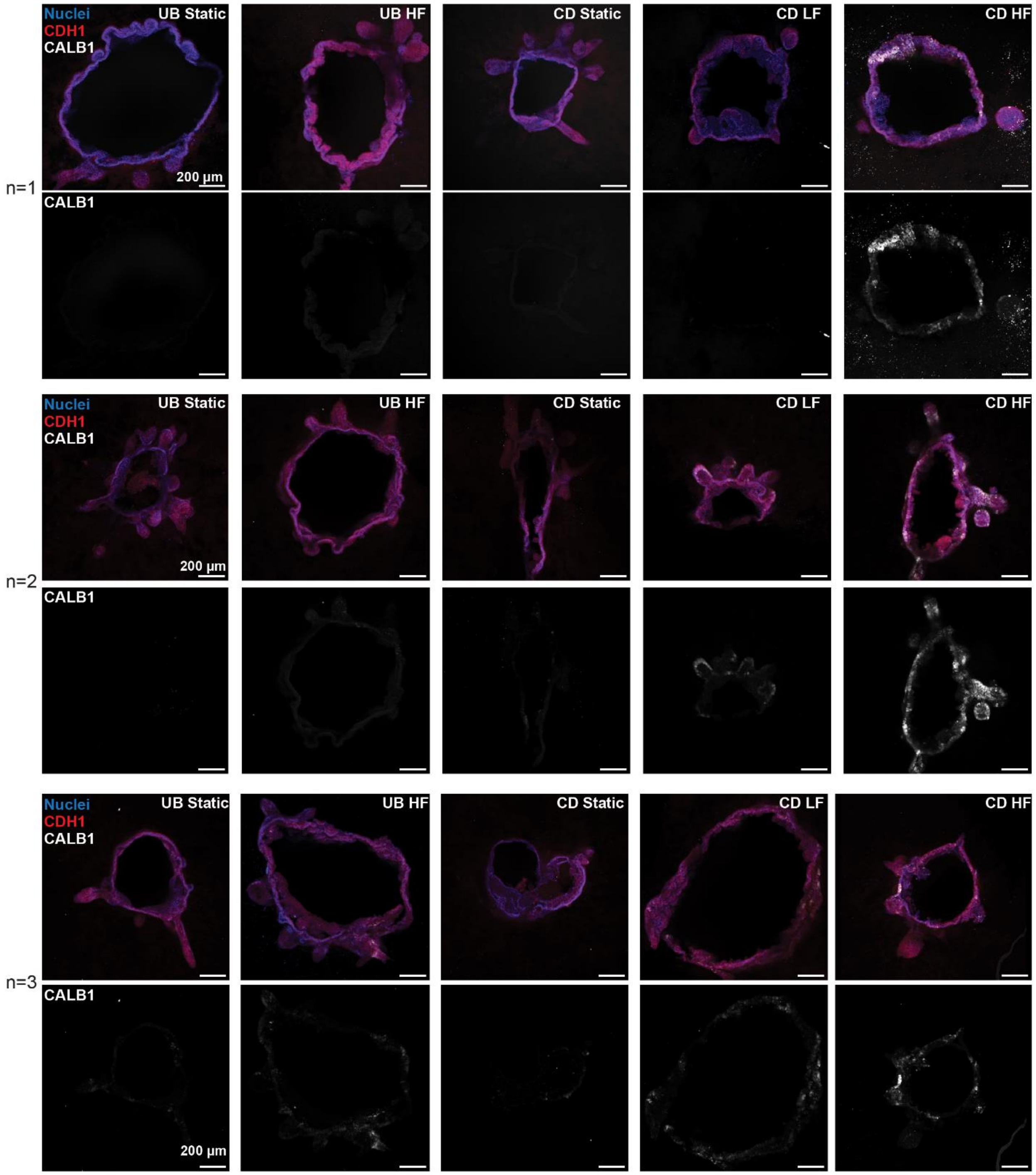


**Supplementary Figure 14. Maximum intensity projection immunofluorescence cross-sectional images of UB and CD on chip**. From left to right, static UB on chip (UB static), high flow (HF) UB on chip, static CD on chip, low flow (LF) CD on chip, and HF CD on chip. Top row: nuclei (blue), CDH1 (red), CALB1 (white), Bottom row: CALB1 staining only. Scale bar = 200 µm. Cells cultured for 14 days after seeding on chip followed by 7 additional days with or without CD medium. N=3 independent experiments.

**Supplemental tables:**

**Table S1.** Resources and Reagents

| REAGENT or RESOURCE | SOURCE | IDENTIFIER |
|---|---|---|
| **Chemicals, Peptides, and Recombinant proteins** | | |
| DMEM/F12, HEPES | FisherScientific | Cat#11-330-032 |
| mTeSR Plus | STEMCELL Technologies | Cat#100-0276 |
| Accumax | Innovative Cell Technologies | Cat#AM-105 |
| Gibco Heat Inactivated Fetal Bovine Serum (BSA) | Thermofisher | Cat#10082147 |
| Matrigel Growth Factor Reduced, LDEV-free | Corning | Cat#356230 |
| Geltrex | Gibco | Cat#A14132-02 |
| antibiotic-antimycotic | Gibco | 15240062 |
| TeloCol®-10 Type I Collagen Solution, 10 mg/ml (Bovine) | Advanced BioMatrix | Cat#5226 |
| 0.25% Trypisin-EDTA | Gibco | Cat#25200-056 |
| Fibrinogen, Bovine | EMD Millipore | 341573 |
| Transglutaminase | Modernist Pantry | 1203-50 |
| Thrombin | Millipore Sigma | T4648-10KU |
| $CaCl_2$ | Alfa Aesar | 89866 |
| Gelatin | Sigma-Aldrich | G1890 |
| DPBS+/+ (with $Ca^{2+}$ or $Mg^{2+}$) | Corning | 21-030-CV |
| DPBS-/- (without $Ca^{2+}$ or $Mg^{2+}$) | Corning | 21-031-CV |
| GDNF | Peprotech | 450-10 |
| FGF2 | PeproTech | #10771-938 |
| FGF7 | Peprotech | 100-19 |
| EGF | Peprotech | AF-100-15 |
| CHIR 99021 | Tocris Bioscience | 4423/10 |
| LDN-193189 | Reagents Direct | #36-F52 |
| A83-01 | Reprocell | 04-0014 |
| JAK inhibitor I | STEMCELL Technologies | #74022 |
| SB202190 | Axon medchem | #1364 |
| TTNPB | Tocris Bioscience | 0761 |
| Y27632 | Tocris Bioscience | 1254 |
| aldosterone | Sigma | A9477 |
| BSA | Roche | 10735078001 |
| R-Spondin-1 | Peprotech | 120-38 |
| B27 minus Vitamin A | Fisher | Cat#12587010 |
| [Arg8]-Vasopressin | Sigma-Aldrich | Cat#V0377-100IU |
| ITS supplement | Gibco | 41400-045 |
| MEM NEAA | Gibco | 11140-050 |
| Glutamax | Gibco | 35050-061 |
| PenStrep | Gibco | 15140-122 |
| ReLeSR | STEMCELL Tehcnologies | 100-0483 |

| | | |
|---|---|---|
| Embryonic Stem Cell Freezing Med. | Millipore | ES-002-F |
| DTT | Roche | 51033324 |
| DNAse | Worthington Biochem | LS002004 |
| DMSO | Sigma-Aldrich | #MX1458-6 |
| Methacrylated Hyaluronic Acid | Nanosoft Polymers | 13204-50 K |
| Poly-D-Lysine | Sigma-Aldrich | Cat#P6407 |
| LAP | Sigma-Aldrich | 900889-1G |
| MycoAlert Mycoplasma Kit | Lonza | LT07-118 |
| RNeasy Plus Mini Kit | Qiagen | 74134 |
| Triton X-100 | Sigma-Aldrich | 9410-OP |
| formalin solution | Sigma-Aldrich | #HT501128 |
| donkey serum | Sigma-Aldrich | #D9663 |
| Anti-Adherence Rinsing Solution | Stemcell Technologies | 07010 |
| Collegenase Type IV (1 mg/mL) | Stemcell Technologies | 9001-12-1 |
| Sodium hydroxide | Thermo Fisher Scientific | M1064621000 |
| **Other** | | |
| Aggrewell™ 800 | STEMCELL Technologies | Cat#34815 |
| Transwell (6-well) Polyester 0.4 µm pore size | Corning | Cat#3450 |
| Transwell (12-well) PET 0.4 µm pore size | Corning | Cat#353180 |
| Ø1 mm stainless steel pin (316 Stainless Steel Tubing, 0.034" OD, 0.004" Wall Thickness) | McMaster-Carr | Cat#89875K361 |
| 48-well glass bottom dishes, No. 1.5 Coverslip, 6 mm diameter, uncoated | MatTek Life Sciences | Cat#P48G-1.5-6-F |
| Silicone tubing 0.50 MM id | Cole Parmer | 06411-60 |
| PharMed BPT 0.25 ID tubing | Ismatec | 95723-12 |
| Glass slides | VWR | 16004-422 |
| Chip Gasket and Lid | Celltreat | 229164 |
| 32 G (0.10 mm I.D.) blunt needle | Nordson EFD, | #7018462 |
| 1.5 inches and an inner diameter of 0.25 mm, blunt needle | Hamilton | 7748-11 |
| 0.12” 304 stainless steel | McMaster-Carr | 8983K93 |
| Biomed Clear Resin | Formlabs | RS-CFG-BMCL-01 |
| 10k Rigid Resin | Formlabs | |
| Microbore silicone tubing (0.50mm ID) | Cole-Parmer | - |
| Nanodrop 1000 spectrophotometer | Thermo Scientific | - |
| Nanostring TagSet-48 | Nanostring | - |
| Nanostring XT TagSet-192 | Nanostring | 1210000616 |
| Custom DNA oligos | ID&T | - |
| Nanostring cartridge | Nanostring | - |
| MycoAlert | Lonza | LT07 |
| Opti-MEM | ThermoFisher Scientific | 31985062 |
| Lipofectamine Stem Cell Reagent | ThermoFisher Scientific | 18324012 |
| Puromycin Dihydrochloride | ThermoFisher Scientific | A1113803 |
| 3 mL syringe | VWR | BD309657 |
| 100 µL glass syringe | Hamilton | 1710 TLL |

**Table S2.** Antibodies and stains.

| Target | Host Species | Manufacturer | Product # | Dilution |
|---|---|---|---|---|
| **Primaries** | | | | |
| KRT8 | Rat | DSHB | TROMA-1 | 1:100 |
| E-Cadherin | Mouse | Abcam | ab1416 | 1:100 |
| ETV5 | Rabbit | Abcam | ab102010 | 1:400 |
| SOX9 | Rabbit | Abcam | ab185230 | 1:300 |
| AQP2 | Rabbit | Abcam | ab62628 | 1:200 |
| Na+K+ATPase | Rabbit | Abcam | ab76020 | 1:200 |
| RET | Goat | R&D technologies | AF1486 | 1:100 |
| CD117 | Mouse | BioLegend | 313204 | 1:300 |
| Acetylated Tubulin | Rabbit | Cell Signaling Technologies | 5335 | 1:200 |
| CALB1 | Rabbit | Abcam | ab108404 | 1:200 |
| **Secondaries** | | | | |
| 488 anti Rat | Donkey | ThermoFisher Scientific | A21208 | 1:400 |
| 546 anti Mouse | Donkey | ThermoFisher Scientific | A10036 | 1:400 |
| 647 anti Rabbit | Donkey | ThermoFisher Scientific | A32795 | 1:300 |
| 647 anti Rat | Donkey | Abcam | AB150155 | 1:200 |
| 555+ anti Rabbit | Donkey | ThermoFisher Scientific | A32794 | 1:300 |
| 488 anti Mouse | Donkey | ThermoFisher Scientific | A21202 | 1:300 |
| 546 anti Goat | Donkey | ThermoFisher Scientific | A11056 | 1:300 |
| **Cell stains** | | | | |
| DAPI (4′,6-diamidino-2-phenylindole) | - | Millipore Sigma | D9542-1MG | 1:1000 From 1 mg/mL in DI wter |
| ActinGreen 488 ReadyProbes | - | ThermoFisher | R37110 | 1:100 |

**Table S3.** Differentiation medium compositions.

| Additives | Reagent Name | Final Concentration |
|---|---|---|
| Base medium (all added to DMEM/F12, HEPES | GlutaMAX (100x) | 1x |
| | MEM NEAA (100x) | 1x |
| | Pen Strep (100x) | 1x |
| | B-27 Supplement minus vitamin A (50x) | 1x |
| | ITS Liquid Media Supplement (100x) | 1x |
| ME Additives (add to base medium) | LDN-193189 | 10 nM |
| | CHIR99021 | 4.5 µM |
| UB-I Additives (add to base medium) | FGF2 | 200 ng/mL |
| | TTNPB | 0.1 µM |
| | LDN-193189 | 30 nM |
| | A83-01 | 0.2 µM |
| UB-II Additives (add to base medium) | FGF2 | 200 ng/mL |
| | TTNPB | 0.1 µM |
| | LDN-193189 | 30 nM |
| UBCM (add to base medium) | LDN-193189 | 200 nM |
| | TTNPB | 0.1 uM |
| | CHIR990021 | 1 µM |
| | JAK inhibitor I | 100 nM |
| | GDNF | 100 ng/mL |
| | A83-01 | 0.2 µM |
| | Rspo1 | 100 ng/mL |
| | FGF7 | 50 ng/mL |
| | SB202190 | 5 µM |
| | EGF | 50 ng/mL |
| CD (add to base medium) | KSR | 3% v/v |
| | Aldosterone | 100 nM |
| | Vasopressin | 1 I.U./mL |

**Table S4.** Fragmented and bulk matrix composition list.

| Abbreviation *(in text abbreviation)* | Fragment Composition | Interstitial Composition | Fraction (Fragment: Interstitial) | Bulk Composition |
|---|---|---|---|---|
| Fragmented matrices: | | | | |
| F1HA_I3.75C_6040 | 1% w/v HA | 3.75 mg/mL Col | 60:40 | 1.5 mg/mL Col 25% v/v BsM |
| F100BsM_I1.5C_6040 *BsM (frag.) - Col I* | 100% v/v BsM | 3.75 mg/mL Col | 60:40 | 1.5 mg/mL Col 60% BsM |
| F50BsM_I1.5C_8020 | 50% v/v BsM | 7.5 mg/mL Col | 80:20 | 1.5 mg/mL Col 40% BsM |
| F50BsM_I50BsM_6040 | 50% v/v BsM | 50% v/v BsM | 60:40 | 50% BsM |
| Bulk matrices: | | | | |
| B_1.5C-60BsM *BsM – Col I* | - | - | - | 1.5 mg/mL Col 60% BsM |
| B_1.5C-40BsM | - | - | - | 1.5 mg/mL Col 40% BsM |
| B_50BsM *BsM* | - | - | - | 50% BsM |

**Table S5.** Gene panel for Nanostring transcriptomic analysis. Table includes gene name, accession number, marker or functional categorization, probe sequences, and references used for categorization. CD = collecting duct, CI = Cilia, DT = distal tubule, EMT = epithelial to mesenchymal transition, GF = growth factor, HK = Housekeeping, IJ = injury, LH = loop of henle, MET = mesenchymal to epithelial transition, MM = metanephric mesenchyme, MS = mechanosensitive, MX = matrix, N = neural, P = proliferation, PO = podocyte, PT = proximal tubule, S = stem cell, SC = stromal cell, UR = ureter, V = vasculature.

| Gene | Category | Accession # | Probe A sequence | Probe B sequence | Reference |
|---|---|---|---|---|---|
| *AADAC* | LH | NM_001086.2 | TATGCTATGAGGATCCCCACAATCAGAAGGTACAGCGATTTTCTTCCCATCAGGTCTCGATCTCGTACAAAACGACTATGACCAT | CGAAAGCCATGACCTCCGATCACTCTCATTCTCCATGGCTCCTCAACGTTATCTGGGAGAGGCGTATAAATATAA | 2 |
| *ABCB1* | PT | NM_000927.3 | TTCTGGCCACCAGAGAGCTGAGTTCCTTTGTCTCCTACTTTAGTGCTATACTTACGGTTTTGACATTCCGGAAATTTGGATCGAA | CGAAAGCCATGACCTCCGATCACTCGCAAAATATGAGGCTGTCTAACAAGGGCACGAGCTATGGCAATGCGTTGT | 3 |
| *ABCC1* | PT | NM_004996.3 | GAAGTCTGGCTGCCAAAACTCCTCCGTGGGAAGCACCAGGAAACCACTTGCAGAAGATCAAAAAACGATCCCTGTCCATCAATAC | CGAAAGCCATGACCTCCGATCACTCCAGTAGGCTGAATTTGGTCACTAGGATCTTCTATACAACCAATTCCTCCA | 4 |
| *ABCC3* | PT | NM_003786.3 | CAGCACAGTTACCCTGTAAGGCACTTTACAGGTGCAAAATCTTAAGGTGCCTAGCCCGCGCAACTCGTCCTATTAAAAGTCGGAA | CGAAAGCCATGACCTCCGATCACTCCTTAGGCGTGTCATTCACCACTTGGGGATCATTTCCTCATCTAAAGCATT | 4 |
| *ABCC4* | PT | NM_001105515.2 | TGCACTGAGAGGATCGTCCAGGAGATAGATGTCAGCATCTTGATACACTGCCTGCATTCTCATGGAAATGCAATGGATTCATTCC | CGAAAGCCATGACCTCCGATCACTCATTTGACAAATACACAGTTCGAACAAGTGTCTGCTAACTTCCGCATCTAC | 4 |
| *ACTA2* | SC | NM_001613.3 | AGCAGGAAAGTGTTTTAGAAGCATTTGCGGTGGACAATGGAAGGCCCGGCCTACGGTTACCGTCTTTATAAGTGAACAAAACCGG | CGAAAGCCATGACCTCCGATCACTCAAGGCATAATTCCACAGGACATTCACAGTTGTGTGCTAGAGACAGAGAGG | 5 |
| *ACTB* | HK | NM_001101.3 | ATCGGCAAAGGCGAGGCTCTGTGCTCGCGGGGCGGACGCGGTCTCGCATTTGGAATGATGTGTACTGGGAATAAGACGACG | CGAAAGCCATGACCTCCGATCACTCCGATATCATCATCCATGGTGAGCTGGCGGCGGGTGTGGACGGGCGGCGG | |
| *AGTR1A* | SC | NM_000685.3 | TTGGAAACTGGACAGAACAATCTGGAACTCTCATCTCCTGTTGCTCCTCTCAGAGTTGTCAAGACGTTGGCGTGATGGGATGAAC | CGAAAGCCATGACCTCCGATCACTCGTAGCACAGTTGCTAATAGCTGAAAACCGGCACGAAAACTTTACTGCCCT | 6 |
| *ALX1* | SC | NM_006982.3 | GGCGCGGGGCAGGGGTCCGAAGGCCTGCACGCATTTGCCCTTGGTGAAAAGTCTCTTGTTATTTTGTGCTCGC | CGAAAGCCATGACCTCCGATCACTCTGCTGTCCTGACAGGGCGAGGTCCTCTCCAAGCGCACGTGATGCTC | 6 |
| *AQP1* | PT | NM_198098.1 | AGAAGGAGGGCCGTGCCCAAAGGCACAAAGCAATTACGGTAGAATCCCAGCACAAGAATCCCTGCTAGCTGAAGGAGGGTCAAAC | CGAAAGCCATGACCTCCGATCACTCTCTTCCCCCTCCAAGCACCATTGGGAGCAAGGTGCATGTTAGGAAA | 7 |
| *AQP2* | CD-DT | NM_000486.5 | AAAAAGGTCTGAGTTCAAGTGTTGGCTCCACCATATACTACCTGTGAGACCAGATAAGGTTGTTATTGTGGAGGATGTTACTACA | CGAAAGCCATGACCTCCGATCACTCTCCCAGGAGCCTAGGATTTGTCTCAGTGGAAAAATCATGGGATTTTGTGT | 8 |
| *AQP3* | CD-DT | NM_004925.3 | TTTCCTCTCCTTATGCACAGATGGACAGGCTGCCTTCCCTTTCGTTGGGACGCTTGAAGCGCAAGTAGAAAAC | CGAAAGCCATGACCTCCGATCACTCCTCCTCCATGTGAAGCCCCTGAAACATACACACCCTGGAAC | 4 |
| *AQP4* | CD | NM_004028.3 | AATCAGGTTTAATAGTGGTCCATTATCTCACTGCCACAGACACACAACTGCCAGCAGACCTGCAATATCAAAGTTATAAGCGCGT | CGAAAGCCATGACCTCCGATCACTCTGACGACTCTCTGGGAATCTCACACATCACTCTTTCCTAGCACCGAAGAG | 9 |
| *ASB4* | SC | NM_016116.3 | GGTCTACAGTTGATTGTAGCATTGTGGTCCAGTAGCACCAGAAGACATT | CGAAAGCCATGACCTCCGATCACTCAATCCACATTGGCCATTTCA | 6 |

| | | | | | |
|---|---|---|---|---|---|
| | | | CCGAGGAAACAAGTGGGAGGTTTGTGAAGAGGTTTA | CAAGCCACGTGAAGAGGGGTTTTCCCATTG | |
| *ATP1A1* | PT | NM_000701.7 | GAGCAAGCCAGGTAGCTGAAGTCTTGTCAAAAGAGACACCACTCTGATTCCAATATGCCAATTGAGGCGAGGGAACTGGGATTGT | CGAAAGCCATGACCTCCGATCACTCCTGGTTAGCCTGAAACACTGCCCTGTTACAAAGACCTGCAATTCTGGACA | 10 |
| *ATP6V1B1* | CD | NM_001692.3 | TCCATGTTCACCCCCATGGCTGCAAAGACGATGGCGAAGTTGTCGCTTACAGATCGTGTGCTCATGACTTCCACAGACGT | CGAAAGCCATGACCTCCGATCACTCCCATGGTTCCATTCTGCTCAAAGTCAGACTTGAAGAATCTGGCTGTC | 4 |
| *ATP6V1B2* | PT-LH-DT | NM_001693.3 | GGGCTCACGTCAGGAAGGCGCAGGAATTCAGAAAACTCTAAAGCTTGGAACACTCACTAATTGTCCGTGCGACGACAAATTACCG | CGAAAGCCATGACCTCCGATCACTCCACAGAGAGACCAACAGGACGCATGGCAGAACTGCATAGAAGATCGCTCA | 11 |
| *AVPR2* | CD | NM_000054.2 | GTGACATAGGTGCGACGGCCCCAGGGCTCCGCAAACCAATTTGGTTTTACTCCCCTCGATTATGCGGAGT | CGAAAGCCATGACCTCCGATCACTCGGGTAGGTGCCACGAACACCATCAGGGCAATCCAG | 12 |
| *B2M* | UB-CD | NM_004048.2 | AGACAAGTCTGAATGCTCCACTTTTTCAATTCTCTCTCCATTCTTCAGTACCACGCGATGACGTTCGTCAAGAGTCGCATAATCT | CGAAAGCCATGACCTCCGATCACTCGGGGTGAATTCAGTGTAGTACAAGAGATAGAAAGACCAGTCCTTGCTGAA | 12 |
| *BMP4* | SC-PT | NM_001202.3 | TCTTCCTCCTCCTCCCCAGACTGAAGCCGGTAAAGATCCTAGCTGTTATGGCTATTGCTGAAACAGCAAAATT | CGAAAGCCATGACCTCCGATCACTCGCGCTCAGGATACTCAAGACCAGTGCTGTGGATCTGC | 13 |
| *BNC2* | PT | NM_017637.5 | ACTCCTCTGTTAGGATGAAAACGAGGTGTTACCACTGGCCTTTTCTATAGCCGAGATACGTCTTACCGCACATAGGTAACCTATT | CGAAAGCCATGACCTCCGATCACTCATGAGTAATGATCAAGAAAGGAACGATCCATGCTGGTGCCATTCGCACCT | 10 |
| *BSND* | DT | NM_057176.2 | CTGAAATCGCTGCAACTCAAGGCTTCAAGGGAGAGCAAGATCCCTTTCAACAGTTTCATGCTAATCAAAGAAGAGCTAATCCGAC | CGAAAGCCATGACCTCCGATCACTCCTGTAGTTCAGCCTAAACACCGAGACAGGCTTACACAGGGAGAGAAGACA | 12 |
| *CALB1* | UB-CD-DT | NM_004929.2 | CCCCAGCACAGAGAATAAGAGCAAGATCCGTTCGGTACAGCTTCCCCTTGACGTAGATTGCTATCAGGTTACGATGACTGC | CGAAAGCCATGACCTCCGATCACTCTAGATACAGTGTATCACTAGCAAGTGGTTGCGGCCACCAACTCTAGTTAT | 14 |
| *CDH1* | UB-CD-DT | NM_004360.2 | CTCTTCTGTCTTCTGAGGCCAGGAGAGGAGTTGGGAAATGTGAGCAATTCCATAAAATTGGTTTTGCCTTTCAGCAATTCAACTT | CGAAAGCCATGACCTCCGATCACTCATGGGCCTTTTTCATTTTCTGGGCAGCTGATGGGAGGAATAACCCAGTCT | 9 |
| *CDH16* | UB-CD-PT-DT | NM_004062.2 | ACATCTGGGCATTGTGGCTGACCACCACGGGGATTATGTGTTCACGCTCGTAACCCCGGGCACTAGATTATGGTGAATGGT | CGAAAGCCATGACCTCCGATCACTCCACGTTGCAGCGACACACGATCACTCGAACCAGGAGCTGCC | 15,16 |
| *CDH2* | PT | NM_001792.3 | TATGATGTCTACCCTGTTCTCAGGAACTTCACCATACTATTTCTGTTCACGGATGAAGGCCTATATCAATG | CGAAAGCCATGACCTCCGATCACTCGGGTTGATCCTTATCGGTCACAGTTAGATTAGCTAC | 7,17 |
| *CDH6* | PT | NM_004932.3 | AGTTGGCTGACCTGACATGGAGTGTAGTAAAACACCAGGACAGCGGAAGCCGGCTATGATTCACCACCGTGTAGGAGTTATGCGC | CGAAAGCCATGACCTCCGATCACTCTTAAAGCACAGTCCTTTATCCCATTAGCCTGTGAAATGTACAGTTAGGGC | 18 |
| *CLCnKB* | DT | NM_001165945.1 | AGAGAGTCTCCCCAAAGAGGCGCCCGATAGCAGCTCCATAGACAAAGATGCGGATGTGTCGCTAACCTCCTGGTCCTATAAAGCA | CGAAAGCCATGACCTCCGATCACTCGATGGGATTGGTGATCCCTCCAGCCACGATGCCCTCAGGGAAGATAAAAG | 12 |
| *CLDN10* | LH | NM_001160100.1 | AACTCAAGAGGCAGCTTGCCAGCGAGCTCTTTTAGACATAAGCATTTTTACAGAAATGTCACTCCCATGGTGGCTGATATAGAAA | CGAAAGCCATGACCTCCGATCACTCATGGGAGGGCCTTGATGGGATCATTTTGTGAACAGTTCGCTTTTATAACA | 4 |
| *CLDN14* | LH | NM_001146077.1 | AAAAATTGCCCAGAAGTAAACTTTGTGCTGGAACCCCTGCCTCCATTGACCATGTCGAACCTTGGATAGGAGCGACCGATTACGT | CGAAAGCCATGACCTCCGATCACTCCCCTGTGCTCTAAAGACATTTCCTCGCATTCACATTATTTCCTTGGATAC | 4 |
| *COL1A1* | MX | NM_000088.3 | GCCCTGCGGCACAAGGGATTGACACGCGTTCCCCAAATCCGATGTTTCT | CGAAAGCCATGACCTCCGATCACTCGTCTTTCAGCAACACAGTTA | |

| | | | | | |
|---|---|---|---|---|---|
| | | | GCTCTGTGAACTGTCATCGGTCCGATCAATTAGTCT | CACAAGGAACAGAACAGTCTCTCCCGCCCA | |
| *COL1A2* | MX | NM_000089.3 | CCTCTCTTTCCTTCTTCACCACTGGGACCAGGAGGACCTTGCCTATGCATCATGTGCCTCACTAGGACATCATGCT | CGAAAGCCATGACCTCCGATCACTCCTGGAGGGCCGGCAGATCCAGCTTCCCCATTAGGG | |
| *COL4A1* | MX | NM_001845.4 | GACCCCTTGGTCACCCTTGTCACCTTTTGGTCCTTGAAAACTTAAGCCCACGGGAAAACCATTGATCATGTATTCTTTGTCGCAT | CGAAAGCCATGACCTCCGATCACTCTCTCCTTTTTCTTGAACTTGAGCTTGTCCTGGTACTCCTGGAGGCCCACT | |
| *COL4A2* | MX | NM_001846.2 | CTCAGTCCAGGAAAGCCCATGATTCCTTCTTCTCCCTTCAAGGAAATGCCCCCTATGTTGTTGTTGAGGAGATTTATGTTGTGGG | CGAAAGCCATGACCTCCGATCACTCCTTCTGTCCTGGTGATCCTTTTTCACCACTCAAGCCAGGGTAACCC | |
| *COL4A3* | MX-DT | NM_000091.4 | CTTTGACCCAGTGGTCCCATTTCACCTTGATGACCTGGAATTCCCTCTAATACATCGTGATACGGGCGATATAATGCTC | CGAAAGCCATGACCTCCGATCACTCCATCTTCACCTGGTGGCCCTAAAATTCCCGGATTTCCTGGATATCCT | 12 |
| *COL4A4* | MX | NM_000092.4 | AGTTCCTTTCTGACCTTTCAATCCATGCAAGCCGTTCAGGCCGAATTTGTCCGCTGGGTAATGGTGGAATTAAAGA | CGAAAGCCATGACCTCCGATCACTCTGGACCAGGTGGCCCCACATCATGCAAACCTGAAGCACCTTT | |
| *COL4A5* | MX | NM_000495.4 | AGAAGCACAGGGAAAAGAATTGTTAATTCTGCGTGTGTGTTTTTGTTTCTCTTTACTTGTTGTCACCCCAGAGATGTTCTCAATT | CGAAAGCCATGACCTCCGATCACTCGTTGTTTCCCAGTTCTGTACAAGCAGAGGCCACAGGAGGATTCTTACATA | |
| *COL4A6* | MX | NM_001847.2 | ACCCTTTTGACCCTCAATTCCTTGGAATCCATTAAGCCCAGGAAATCCCACTCGTCAATGATCTACGATCCTCGCTAAGTATTCA | CGAAAGCCATGACCTCCGATCACTCACAGCACCATCTATATCGATGAAAACATCTGGGCCTGGCAGGCCAATGTC | |
| *CRABP1* | N | NM_004378.2 | CTGGTGCAGACCACGTCATCGGCGCCAAACGTCAGGATAAGTTCATCGCTTCAGTTAAAGGCTATCTTGCTCCGCTCGTTCTC | CGAAAGCCATGACCTCCGATCACTCCTTCCTGAAAGTAGGAGCAAGCCAGCTGCCTTCACTCTCGGACATAAATT | 19 |
| *CUBN* | PT | NM_001081.3 | GAGACGAATCTCAGCGTCAGGGCGCTGCTGAAGGATGTGATAGGATGGGGCCACCGTTTTGCCAGTTCCACCAGTAACTGTAAGG | CGAAAGCCATGACCTCCGATCACTCCTGATGCGGTGACCGTGGTGTGGAAACCCCCAGCACTGATGCTAGAATCA | 5 |
| *DBA* | CD-DT | NM_001022.3 | GCTTCGTTTGCGTCGTTCGTGGCTGGAAGGAACAGTGGTGGAGAATACTACCGGCACAAGCAGACAAAATCAACATGGTCATTTA | CGAAAGCCATGACCTCCGATCACTCGTGTAATAGAGAGAGGATAGAGAGCTCCTGTTCGGAGCTGGGGGAACTTG | 2 |
| *DUSP9* | LH | NM_001395.2 | CCCCCACCCCCATCCCGCAAAGAAAAAGTAACAGTCGCAGAAGAGGCAGGCAGTTCTTAATTCCATTGGCTCTCTCAATGGGTCG | CGAAAGCCATGACCTCCGATCACTCCCGGGACATGGGCCTGGACAACCACCTGGAGAGGGAA | 12 |
| *EBF1* | SC | NM_024007.4 | AGTCTCAAAGGAGATTCTCACACAGTATGTGTGCCACCAAGCTCTCGAGACAGTGCGTTTTATAAATACAGGGATGCTGTTTTTC | CGAAAGCCATGACCTCCGATCACTCCAGAGTGTGGAATTCTGTGCCTGGACCCTGTTCCATGA | 6 |
| *EGF* | LH-DT-GF | NM_001963.4 | TATTTTCTCTGATGTCTCCAACAGAGCCTTCACTCCCGATATTTTGCTGATGTAACTGTTACTGACCTTGC | CGAAAGCCATGACCTCCGATCACTCCAGCCGCTTATCAAGCACATCCAATGACACAGCTGT | 20 |
| *ELF3* | CD | NM_004433.4 | GAGCCCAAGCACTGAGCAGGGAGGTGAACGCCAGTCCAGAAAGAAGCTGTGTCCGTCTATACGCATACTGGTCCACATATA | CGAAAGCCATGACCTCCGATCACTCACATATTTATATAGCACATAAATTAGGGAGTGCTCTGACCCCTGCCCGTG | 2 |
| *EMX2* | UB | NM_004098.3 | AGAAGTTTGCATCAGGGTTTTAAGCCTCAGGATGTTAGGAAAGGGAATGTCACCGTGTGGACGGCAACTCAGAGATAACGCATAT | CGAAAGCCATGACCTCCGATCACTCGCATTGCAGTGAATGGTTGAACTCGGCCAATTTCTCCAACCACTGAAAGG | 9 |
| *ENO2* | N | NM_001975.2 | GCAAAGCACAAGTGACACATGGTCCCTCTCTAACACCTCAGCACACCAACCCCGAATGTATAATGCTGACGTTCTTGCTTTTGGC | CGAAAGCCATGACCTCCGATCACTCTATGCACAGTTCACGGCTCATATGCAAAGTGGAAGACACGTGGGACAAGA | 21 |

| | | | | | |
|---|---|---|---|---|---|
| *ETV5* | UB | NM_004454.2 | TTCCATCAGGTTGTCTGGTGCAGAAAGCACAAAGCAGGCAACAAGGCGGGCCTCAAGACCTAAGCGACAGCGTGACCTTGTTTCA | CGAAAGCCATGACCTCCGATCACTCTGTCCAGGCTGCCCTGCACACTTCAAAAATGTACAACTCAGGTGCAAATG | 9 |
| *FGB* | MX | NM_005141.5 | ATTGCTATTCACAGTCTCATCTATATATAATTGGTGCTTTTCCAGTTCTGCGCTTTAGAATGCTTAAAGATGGCAGAGTTGGAGG | CGAAAGCCATGACCTCCGATCACTCCTTCTCAGGTTTTCCAGGATTGAACGAAGCACACGAAGGTTAGTTGGGAT | |
| *FGF8* | MET-DT | NM_033163.3 | CCAGCCTGCAGAGCAGCGCACAGCTCATCTTGCTTGAGCTGATGTACAGGGTGAAGGAACTGTAACGGCAGCT | CGAAAGCCATGACCTCCGATCACTCAACAGCAAACAATATCAACAACCGGAACCCAGGGCTCCCCCAGCACCTCC | 12,18 |
| *FN1* | MX | NM_212482.2 | TCGATGTGGTCTGCACAGAGGTGTGCCTCTCACACCTCCCCTTTCCCAAGTAAATGTACGGGAATTATCG | CGAAAGCCATGACCTCCGATCACTCCTGCGGTTGGTAAACAGCTGCACGAACATCGGTGAAGGGGCCAGATCCGC | |
| *FOXD1* | SC | NM_004472.2 | GCGCCTGGAGGAGCGAACAAAACACCGAACCACCAAGACGAGAAAAGGAGCACAAACTCACTACTACCAACAACCTCACCAAAAA | CGAAAGCCATGACCTCCGATCACTCTCCGAGAATTCGCAGCGGCGAAAATGGGCGCGCGAGGTCGAGAGGGGCC | 6 |
| *FOXI1* | CD | NM_005250.2 | AAGCGTTGAAAGGCGGACGGAGGCTGGAGGAGGCGGCCAGGAGCGCCTGCCAATGCACTCGATCTTGTCATTTTTTTGCG | CGAAAGCCATGACCTCCGATCACTCCGAGCTGGTAAAAGCCGCCCTGGACATGCGGGTCGAGCATCAGGG | 9 |
| *FOXQ1* | CD-DT | NM_033260.3 | ATGTCTCTCACACTCAGTCATACCTGGAAATGCCACATACGTACACGGATCTCTAAGGTAAAGTGCTTCTCAATAACATCCGCTC | CGAAAGCCATGACCTCCGATCACTCGCACAGTGAAGCATCGGTCAATAGCAAGAAATCACTGAAGAGCCTCTGAC | 12 |
| *FXYD2* | DT-LH | NM_021603.3 | AGGCCCCCATTGCGAACGGTCTCATAGTCATAGTAGAACGGGTCCCAATCAGAAACCCTACTTGCCTGCTGCATGTTGTG | CGAAAGCCATGACCTCCGATCACTCAGGAGGATGAGGAGCCCCACGATGAAGGCCAGTCCAGCGAAGATC | 12 |
| *GAPDH* | HK | NM_001256799.1 | TCCCCTCTTCAAGGGGTCTACATGGCAACTGTGAGGAGGCTAGGACGCAAATCACTTGAAGAAGTGAAAGCGAG | CGAAAGCCATGACCTCCGATCACTCTACTTTATTGATGGTACATGACAAGGTGCGGCTCCCTAGGCCCC | |
| *GATA2* | SC | NM_001145661.1 | CGACTGCCCGCCCATATTGCACTTGGTCACTACATCTATTTGTGGCAGAATATGTCGATGCAAAAAACCT | CGAAAGCCATGACCTCCGATCACTCGACACTAGTACAAAATAAGTTACTTCTGGCCGTGGTGCTCCCTGCAG | 6 |
| *GATA3* | UB-CD-DT | NM_001002295.1 | TTTCCTTTGCAGAGGCCAACAGCTTTGAACAAATGATTCGCCTACAACTTCTTTCGGGTTATATCTATCATTTACTTGACACCCT | CGAAAGCCATGACCTCCGATCACTCACCCTCAACGGCAACTGGTGAACGGTAACACTGATTGCCCAGAACTGGTA | 9 |
| *GDNF* | GF-SC | NM_000514.3 | AAGTTTTTCAATATTTTGTCGTACGTTGTCTCAGCTGCATCGCAAGAGCCCGCTTTATTATGTGTTCGTCTAACTCTGTTTCTGT | CGAAAGCCATGACCTCCGATCACTCCCTACTTTGTCACTCACCAGCCTTCTATTTCTGGAT | 22 |
| *GFRA1* | UB | NM_005264.4 | GCACACGGTGGCAAAACATGAGTGGGAATTTCATTCTCAGACCCTGCTGGCTTCCTTCCTGTGTTCCAGCTACAAACTTAGAAAC | CGAAAGCCATGACCTCCGATCACTCTGTGTATTGCCCGACACATTGGATTTCAGCTTCTGTGCCTGTAAATTT | 9 |
| *GJA1* | DT | NM_000165.4 | TTGCCTGGGCACCACTCTTTTGCTTAAAAGTAGTGAAGTCACGCCAAGTGCTAGCATCTCTGACGAAAACAGCAGACGGAAAAGT | CGAAAGCCATGACCTCCGATCACTCTTGTCAAGGAGTTTGCCTAAGGCGCTCCAGTCACCCATG | 12 |
| *GPD1* | LH | NM_005276.2 | ATCTGAGCCTGTTATGAAAGGTGTAGCTCTGTCAGCAGATTAAGTTCCTGCATGAACGTGTCGTGTTATGCAGCGGTATGTCGTG | CGAAAGCCATGACCTCCGATCACTCCTACTGAGAAGGAAGCCTGGGTGAATACAAAGGTCACAGGGAGCTGAGTG | 2 |
| *H2AFX* | IJ | NM_002105.2 | CTAGGTGCTTGGATTGCCGAGTTGAGTTTGCTGGAAGGGAAATGGGGCGGCCTAAATTGGGAAAAAAGGTTTTAGCTATTGATGG | CGAAAGCCATGACCTCCGATCACTCCAACGGAGGCTCAGTCCAGACAGGGATTAACCGACTTGTGCTGGTAT | 23 |

| | | | | | |
|---|---|---|---|---|---|
| *HAVCR1* | IJ | NM_001099414.1 | TGATTGTTATTCCAAAGGCCATCTGAAGACTCTGTCACGGTGTCATTCCCCCTACATATATAGGAAAAGGGAAGGTAGAAGAGCT | CGAAAGCCATGACCTCCGATCACTCCTTTAGTGGTATTGGCCGTCAGTAGACTATGTTCTAGGAACAGTTGAGTT | 24 |
| *HGF* | GF | NM_000601.4 | GCAGTATTCACTTTTTTGGTTTTTATCTTCAGTGCTGGATCTATTTTGATCCTGTTGCAGTATCACGTAAATACCTACTTCGATA | CGAAAGCCATGACCTCCGATCACTCAAGTGAATGGAAGTCCTTTATTCCTAGTACATCTATTAGCACATTGGTCT | 25 |
| *HIC1* | SC | NM_006497.4 | CGACTCCAGAACTCCCCGCCCTTGGTGACTGAGGAAGGAAGGGTTGGCCCCTCAGCATCTTTAGCAGTAGCACTTGCTAAATTGG | CGAAAGCCATGACCTCCGATCACTCCGCGGGAGTGGGAGACCTGGTGGTAGGCTCTTCGCCTTC | 6 |
| *HNF1B* | UB-CD | NM_000458.2 | TGCATAGAAGGGAAACTGGGCTTCGGGCTGCGCCTCCTGGTCAAGACTTGCATGAGGACCCGCAAATTCCT | CGAAAGCCATGACCTCCGATCACTCAGACAGGAGTCCTTGACATCGTGGGAGAGGCATTGTGGCAATAC | 9,26 |
| *HNF4A* | PT | NM_178849.1 | GAAGCTATTCAGCCACTGTAGTTAAGAGCTCCTGTTCTGATCCAGCGATTAAGCCGTAGTTGAATTTATGGAGCGGTGCC | CGAAAGCCATGACCTCCGATCACTCAAGCCCCTCAACTTGATCACGGTGAGAACACAGGGAGCCTTTGGA | 27 |
| *HOPX* | SC | NM_001145460.1 | CCAGTGAAGCCACTGCTTTACACAGAAGATACACATAGCTTCCTATTGTTCATTTGGTGTTGCCTAGTACTAGGTGACTGGTACC | CGAAAGCCATGACCTCCGATCACTCACACCATCTGTCATCATGACTCAAAGGGAAATGCTAGCCACACCATTTTT | 5 |
| *HOXA11* | UB-DT | NM_005523.5 | TCTTTTCCCACAAGGATTTGCAGCCCTCTTCCACCTCAAAGCTACCTCCACCATGCTGAAATCATGGTTGAAGAAGAAGAAGATC | CGAAAGCCATGACCTCCGATCACTCGGTGGCTAGGAGGAGTGGAAAGACACTGATGCCACCTGGAATCATAGATT | 12,18 |
| *HOXA13* | SC | NM_000522.5 | ATCAAAACCTGCCCTCTCAGACACATGCAGACCCAACAGCAGATTTTAAACATCAACACCAAACGCAATATCAGGCTGAATGGTG | CGAAAGCCATGACCTCCGATCACTCCGTCAAGGTCAAGGTTTAAGGCCTTTCGTAGTCCTGTCATTTCAACAAAT | 6 |
| *HOXB7* | UB | NM_004502.3 | AGGGTTAGTCCAGACCCACAGTCCACAAAGACAGCATTAAAGAGGGCTTACCTAGTTGACTTGGAGTATGCCATGAAGACTCGTC | CGAAAGCCATGACCTCCGATCACTCTGGGAAGGCGGAGGCTCAGGAGGGCAGGCAACCAC | 12 |
| *IL1RN* | UB | NM_000577.3 | TCAGCTTCCATCGCTGTGCAGAGGAACCAACCGGGGCGAGCTATTGTAAATACGCAGCAACAAGCGTCCAG | CGAAAGCCATGACCTCCGATCACTCCTTCGTCAGGCATATTGGTGAGGCTGACGGGCTGG | 12 |
| *ISL1* | SC | NM_002202.2 | CAAACTTCTCTGCTCCAACTCAGCTCCATCGCCATTGGTCTGACGCAGCGCCTCGAACAGTGATACGCACACTGATAACTATGCG | CGAAAGCCATGACCTCCGATCACTCCGGCAGACTCGGCGCGGCTCGGGCACCTCTCTTCCTTATCTCTTACT | 6 |
| *ITGA8* | SC | NM_003638.2 | GCTCCTCAATGAAGTTCCGCAGCTGTAGCCAAATGGCTCCCTGTTTCATTAATGTAACGGACAGGTCGCTGAGGT | CGAAAGCCATGACCTCCGATCACTCCAGGAAGAACATGAACTGCACCAGCGTGCTCGTCCATGTGCCTTTCATTT | 6 |
| *JAG1* | GF-MM | NM_000214.2 | CATCCCTCTGTCAGGCAGAGCCTATTATACCCCGATTACCAGAACAGCTCTAACCTGCATACATATGGCATTTAGTTGTTCA | CGAAAGCCATGACCTCCGATCACTCTGAGAACTCCAGATGAGTTTTGTGGTTGAAAAGCCTTTCAGTTCTTCCTC | 8 |
| *KCNJ1* | DT-LH | NM_153766.1 | CCTTTGTCTTGGATACTATGGGAGCAAAACGGTAGCCCCAAAGCACCTCCCGATACCCGTAGCGTGATGGTCTTATAGCTGCTCT | CGAAAGCCATGACCTCCGATCACTCCACTTCCACTGTCTTGCTAAAGTTATGGAAATCCACTCGGTATTTCCCTT | 12 |
| *KDR* | V | NM_002253.2 | CACTCACTTCCATAATCGTCAGTACATGCCCCGCTTTAATTGTGTGATTGCATACGAAATTTGAGCAAGCAATTGAAGGCTTAGA | CGAAAGCCATGACCTCCGATCACTCCTTTGAAATGGGATTGGTAAGGATGACAGTGTAATTTCCTGTGTCTCTTT | 10 |
| *Ki67* | P | NM_002417.2 | CTGATGGCATTAGATTCCTGCACGCTAAGAGTTCTCCCTCTACATCTGCTGTAACGGTAGATGGGAAAAAGTGTGCTCATTTT | CGAAAGCCATGACCTCCGATCACTCGTCTTTCTCTTCACCTACTGATGGTTTAGGCGTGTGCATGGCTTTGCCTG | 28 |
| *KLF4* | S | NM_004235.4 | TTTGGATTCCTCATTTTTCCTGATTATCCACTCACAAGATGACTCAGTTG | CGAAAGCCATGACCTCCGATCACTCAATGATAGAAGATCCAGTC | 29 |

| | | | | | |
|---|---|---|---|---|---|
| | | | CCTACGTATATATCCAAGTGGTTAT GTCCGACGGC | ACAGACCCCATCTGTTCTTTGAT TTTTGTCT | |
| *KLF5* | CD-IJ | NM_001730 .5 | AATTGCCAGTTTAGAAGCAATTGTA GCAGCATAGGATGGAGGTGGGGTT ACATGTTGGAGTTAACGGAGACCC GCCATCGTTTAC | CGAAAGCCATGACCTCCGATCA CTCTGGATGTTTTGTGAGTTAAC TGGCAGGGTGGTGGGTAAATTT GGATTGTG | 30 |
| *KRT18* | CD-DT | NM_199187 .1 | GGCCAACCGGGCATGGACACGGA CAGCAGGTGTTGTTGCGGTTGTTA ATATGACAGGCCGCTAAAGACGTT CT | CGAAAGCCATGACCTCCGATCA CTCTGGAGCGAGTGGTGAAGCT CATGCCCCCAGAAACGGGGT | 12,14 |
| *KRT5* | UR | NM_000424 .2 | CTTGACTGGCGAGACATGGTGGCT TGTTCCTGGTGGAGCAAGAGAACC AGCCTTCTTGAAGACCTATGTAAAG AAACGGGTCACT | CGAAAGCCATGACCTCCGATCA CTCTGAAGCTACGACTGCCCCC GCTCCGGAAGGACACA | 31 |
| *KRT8* | UB-CD | NM_002273 .3 | CTTCCCATCACGTGTCTCGATCTTC TTCACAACCACGGCCCGAACCTAA CTCCTCGCTACATTCCTATTGTTTT C | CGAAAGCCATGACCTCCGATCA CTCCTGTTCACTTGGGCAGGAC GTCAGAGGACTCAGACACCAG | 9 |
| *LAMA1* | MX | NM_005559 .2 | AATAAATCTTCAGCAGCCTTGAGTT CAAGGGTGGCATTTTGGTGCAACT GCAATCTATATTCCTTACACGCTTG CTTGCTCCTAC | CGAAAGCCATGACCTCCGATCA CTCATACCTCCAATTCTTCCAGC GGCTTCTGGTAATTTTCCTGAAT TTGTGAC | |
| *LAMA2* | MX | NM_000426 .3 | CTGTGCTGATGCTCAAGTCATTACC CTCTAAGATAATCATAAGCTGGAGA CTTTGACAGAGAGTCAAAATCAATA AACTCCGATC | CGAAAGCCATGACCTCCGATCA CTCCAATACATTAGTATGTTCTT CAGATGGGTGCAGGTACACCTC ATCTTGGG | |
| *LAMA4* | MX | NM_001105 209.1 | CTCAAAGCCATTTCTCCGCTGACAT CCAGTAGTGCTCTTCCAATTAAAGA ACATGCGTAAAACGACCTCGTTGT A | CGAAAGCCATGACCTCCGATCA CTCAGAGGAGCCACAGAGGCA GAACCGAGCGCCAGGCTGAG | |
| *LAMA5* | MX | NM_005560 .4 | GAGGGAAGCAGCAGCGTTTCGGC AGAACAGGGATGAGCTGCTGCCG GTGCCTGACTAATCGTGTAGGTGT CTAATACA | CGAAAGCCATGACCTCCGATCA CTCCGTGGCAGCCACATGGAC GGGCTCCGTTGTTATAGAAGAG GGA | |
| *LAMB1* | MX | NM_002291 .3 | GCATGAGCACTGGCCTGACTCCGC AAAGCAACTGTTGTTTAAGGCTCC GCCGCGTTTCGTTGCGATAACCTC ATATAGTCCC | CGAAAGCCATGACCTCCGATCA CTCTAGTAACCAGGTTCCACTT CGTTGCACTGACGTCCAATCAT GTGAGGCCG | |
| *LAMB2* | MX | NM_002292 .3 | AAATATCAAGTACCTGGCATGTGGT TGCAGAGTCCCTTGGATGCGATCA TCGAGCTTGGTTCTGTCCCTATAG CTACAGCTGTCT | CGAAAGCCATGACCTCCGATCA CTCTTCAGATGCAGCTTGTAGG AGATACCAGGCTCAAGGCAGAC AGGATTAGG | |
| *LAMC1* | MX | NM_002293 .3 | AACTTCACCCAAACTAGCTTGGCA GTCAATCTTCAGTCCTCTTCTGCAA CCCGTAGGAGTCGGTTGCCACTCG TTGATAGTCAAC | CGAAAGCCATGACCTCCGATCA CTCAAGGAAACCAAACCAAACC ACCCCATTGTGGCCTGAGACTT GCTGGAGTG | |
| *LAMC2* | UB-MX | NM_005562 .2 | TGTGCCGGTAAAAGCCATTCTTGC ACTTCTCGCAGTGAATGCCATCAG TGCTTACTCTTGCCAGTGGGCATG AAATTGCGAGCTT | CGAAAGCCATGACCTCCGATCA CTCAAGAGAACCTTTGGAGTTA CAATTGCAGGGCAAACAGCGGT CCCTTTCTC | 32 |
| *LAMC3* | MX | NM_006059 .3 | CGTAAGCTGCTGACTGGATGGATT AGACTTGAACACGTGTGGGTCATA CACTAATGATCCCGCGGTGTAGTA ATTCTACGGAATG | CGAAAGCCATGACCTCCGATCA CTCCACACAGGTGGCTCTCACA ACAGCTGTGGACTGTAATGTGT GTGGAC | |
| *LHX1* | MET | NM_005568 .3 | ACACACTCGGCCGTGCAAGCCAAA ACTCGCACCAGGAAATCAAAGTAG GGCTGAGGCTGTTAAAGCTGTAGC AACTCTTCCACGA | CGAAAGCCATGACCTCCGATCA CTCACTCCTGAAGACAATCGGC ACGAGCTCCCCAGTCTCTCCAA AAAGAGGAC | 18 |
| *LRP2* | PT | NM_004525 .2 | AGCACCAAACCTAGAGCCCTCCCC TCGCACAGTGTAATACACAACCAG CTCGGATTGGATGTCTCGTAAGGA ATACTTAT | CGAAAGCCATGACCTCCGATCA CTCACAAGATTATTGCGGCCGG ATTCAAAGTTGGGGATGTAGGC ACGTTTGAT | 10 |
| *MAP2* | N | NM_002374 .3 | GGTTTCTTGTATGGCAGTTACACTG GTTGCCATATCCACTGGGTCACCA | CGAAAGCCATGACCTCCGATCA CTCCCTGTGTAAAATCTCTTAGC | 19 |

| | | | | | |
|---|---|---|---|---|---|
| | | | ACTTAAAGCTATCCACGAATGTCAAAAATGTGGTTT | TGAGAATGAGTGGTGTGGGTTTGCTCCTAG | |
| *MCAM* | V | NM_006500.2 | AAAATGTCACAGGAGACTTTGAAGAGGCAGACAGGAGGGTGGTGGCAGCACTATCAATTCGTGACCCCGATCATCCAGTCCAGAA | CGAAAGCCATGACCTCCGATCACTCGCACGTATCTTTTAAGGAATGACACCAGTTCCTGGCTTCTGACCAAAGAA | 10 |
| *MEIS1* | SC | NM_002398.2 | CCTATGAAGAATTAGGGATGCTCCTTGGTCCAGAGTAGATGCCAAGAATGCATTCGCAACCATGTGAAGTAATGTGAGCGTACTT | CGAAAGCCATGACCTCCGATCACTCGCATGTCCCCCAAATTGACTTCAGTTGGTATTTCCTGCTTTTTAAAGGTC | 5 |
| *MEIS2* | SC | NM_002399.2 | AGAGACCAGCTTAACTCAGAGCACACTCCCTCCAAGCGGTGCACCAGTTAGCGTGGCGTATACCATGTTGTTAACA | CGAAAGCCATGACCTCCGATCACTCCTGCGCAGATCCAGCAACCTTGTCAATACTCATTTTTTTCCAGGACCAGA | 5 |
| *MMP1* | IJ | NM_002421.2 | AGAGTCCAAGAGAATGGCCGAGTTCATGAGCCGCAACACGATGTAAGTTGCTATGATAAATGGTTTTCATCAGCCGGATTTTGTT | CGAAAGCCATGACCTCCGATCACTCACTGAAGGTGTAGCTAGGGTACATCAAAGCCCCGATATCAGTAGAATGGG | 33 |
| *MMP2* | UB-IJ | NM_004530.2 | AATGCTGATTAGCTGTAGAGCTGAAGGCACGGCTGCCAGGCAGCAAGAAGGAGTATGGAACTTATAGCAAGAGAG | CGAAAGCCATGACCTCCGATCACTCCACCGGGAGGAGCCACTCTCTGGAATCTTAAATTACCAGGTAGGAGTGAG | 12,33 |
| *MMP7* | UB-CD-DT-IJ | NM_002423.3 | GCACTCCACATCTGGGCTTCTGCATTATTTCTATGACGCGGGAGTTTAACCTATCAGCTAATAGGGTCGGCTCAACAGTGTATCC | CGAAAGCCATGACCTCCGATCACTCGGAAGTCCATTTTGGGCTATTTGGAAATAGTGAGTATTCTGCAACATCTG | 12,33 |
| *MUC6* | UB | NM_005961.2 | GGAGATGCAGACACTGATGCAGTCGTGGGATGAGTGGACAATGAGGAGTGCTCAAGAGCTTGCGAACAAGACGGAAGAAGATATG | CGAAAGCCATGACCTCCGATCACTCGGAGAGTGGCCCTAATGGTAGTAGAGGCAGCTGGAGAAGAAGGAAAAAGA | 12 |
| *MYC* | S | NM_002467.3 | CGCCCGCTGCTATGGGCAAAGTTTCGTGGATGCGGCAAGGGTTGCGCTATGCAGACGAGCTGGCAGAGGAGAGAAATCA | CGAAAGCCATGACCTCCGATCACTCCGTCCTTGCTCGGGTGTTGTAAGTTCCAGTGCAAAGTGCC | 29 |
| *MYH11* | SC | NM_001040113.1 | AATGTCCTCTCGTCTCTGGCTTGGCGAATTGCCCGTGATTTTTCTAGCAGCGTTGGGAATGCCATCTCTATCCCTTCTTTTTCAA | CGAAAGCCATGACCTCCGATCACTCCACTTCTCATCTTCTCCTTGGCTCCAGCAATCATGTAGTAAAAGATGTGG | 6 |
| *NANOG* | S | NM_024865.2 | TCAGTTTCACTCATCTTCACACGTCTTCAGGTTGCATGTTCATGGAGTAGCACCCCTCCAAACGCATTCTTATTGGCAAATGGAA | CGAAAGCCATGACCTCCGATCACTCGGAGGGGAGAGGAAGGATTCAGCCAGTGTCCAGACTGAAATTGAGTAATA | 29 |
| *NGAL* | IJ | NM_005564.3 | CTGATCCAGTAGTCACACTTCTTTTTCCTAAACAGGACGGAGGTGACATTCACGCAGCTCTTAGCGCAATGTATGTGCTGTTATC | CGAAAGCCATGACCTCCGATCACTCTAATGTTGCCCAGCGTGAACTCGCCGGGCTGGCAACCTGGAACAAAAGTC | 24 |
| *NPAS2* | PT | NM_002518.3 | TATTAAGGCCCGAGGCACCCACGTCGAGTGTGTTAAAGTGCTGCCGAGGTCGCTGGGTTAAATAGAGAAATGGCTACTCATTTAG | CGAAAGCCATGACCTCCGATCACTCGTGCGAGGATTTGTGGGAACTTCTTGAGGACGCCGATGGCGAATGACTGG | 10 |
| *NPHP1* | CI | NM_001128179.1 | AAATAGACAGAGGCGTACATGTCTGCTGAGAACCTGTATGCTCATTCCTGCCTTACGACTTCACTGCAATTGACGATTCAGTTAA | CGAAAGCCATGACCTCCGATCACTCTGCCATGTGGCTCTGACTGTATGAATGTTGCTCAGAACCTTATTACCATC | 10. |
| *NPHP6* | CI | NM_025114.4 | CTGTAATCCACTGGTTAGTCTTTTAATTTGCCTTTTCAGTTCATCATTCTCCTCATACCAATGTAAAGTATAGTTAACGCCCTGT | CGAAAGCCATGACCTCCGATCACTCCTTTGGAGTTCTTCAATTAGACTTTGTTTATTATCTGTCAGGGGTTTGCC | 10 |
| *NPHS1* | PO | NM_004646.3 | AAGTTTAGTTTCTAAAGCCAACCCCAGGATGCACCTTGTTTCTACTCTGGCGTGAACCAGATTATGTATGGACGCGCAATAGATA | CGAAAGCCATGACCTCCGATCACTCAGCATTTCATTTTTGAGACGACGTTTACAATCTGCCCTGTCCTTTTGGAG | 7 |
| *NRK* | LH | NM_198465.1 | CAGAACCCTGGATTGCAAAGAGCGCACTTCAATGGCCTTTTGGCCCCAT | CGAAAGCCATGACCTCCGATCACTCGTGCACAGGAATCTCAGCT | 12 |

| | | | | | |
|---|---|---|---|---|---|
| | | | CCCCCAACAACCGCCGGATATACGTTGGGATATAAA | TCTTAATTGACCTGCGCTTCAGCTCACTTTC | |
| *OCT4* | S | NM_002701.4 | TGCCCCCACCCTTTGTGTTCCCAATTCCTTCCTTAGTGAATGAAGAACTTCTGACACATTAGTAACGTCGGCAAGCACTTAGTCG | CGAAAGCCATGACCTCCGATCACTCCATCATTGAACTTCACCTTCCCTCCAACCAGTTGCCCCAAACTCCCC | 29 |
| *ODC1* | IJ | NM_001287189.1 | CCACGCAGGCCCTGACATCACATAGTAGATCGTCGGCCCCCTCTTAACTATAGCTCCATTTGGACAGACGTT | CGAAAGCCATGACCTCCGATCACTCCTACTTCGGGTGGGAAGTCGGGGTTCTGGAATTGCTGCATGAGTTG | |
| *OSR1* | SC | NM_145260.3 | CTCACTCTCCGCAGTCAGAGTTACAAGCTTATATGGTCAAAGACAAAAATCGCGATACTGATGATGAAATTTTCAAAAATGGCTG | CGAAAGCCATGACCTCCGATCACTCCTGCCAGTGCTAAGGACTTCGTTGCCTCTCCTCTCTCCA | 6 |
| *PAX2* | MET-MM-UB-CD | NM_000278.3 | AAATAAATGTAGACTTTTTGTGGTTCTGTTCAGTTCCTGCCGGAGGAGGACAACAGCCACTTTTTTTCCAAATTTTGCAAGAGCC | CGAAAGCCATGACCTCCGATCACTCCTGGTGGGCTTTTGTGTTGTTTTGTTCCTTTTTGCAAAGACCATCATATT | 7,9,34,35 |
| *PAX8* | MET-PT-LH-DT-UB-CD | NM_013953.3 | AGAGGGAAGTTCTGCCATTGAAAAACCAGCCGCATCTCAGATCCCTTCAGCACTACGATCCCATGAGAACACCTGTAGATACTCA | CGAAAGCCATGACCTCCGATCACTCTCAGGAAGACTCTCAGATGTCATGGATTCGGAGTCGCGCTGC | 18 |
| *PCBD1* | DT | NM_000281.2 | CTGCTTGAAGATGGCATCACGGCCTTCCAGCTCATTCCTGGTCTAGGTATCTAATTCGTGGGTCGGGTACT | CGAAAGCCATGACCTCCGATCACTCGCCACTCTTGTCATGAACCCAAAGGCCCTGTTGAAGTCTTTGAAATGAAA | 4 |
| *PDGFRA* | SC | NM_001347829.1 | GAGTTAACAGTTCAAAAGTATAGCTCTTCACAGCATCTTCATTTTGAGCTCGTGGTATCCCTGGTTGAGACATTGGACTAGTGTA | CGAAAGCCATGACCTCCGATCACTCAGTTGAGCCATGGTGATCATCGACCAAGTCCAGAATGGATGAAGGAACTT | 6 |
| *PDGFRB* | SC | NM_002609.3 | CTGGTGCAGGCTCCTGAAGGCTCAGGAGAACAGAGGGATGCCTGAATCAATAGAACAATATCAGTTATGGCGGTG | CGAAAGCCATGACCTCCGATCACTCGTCCCAGAGTGGGTAACAGCTGAGTAGAAGGACAGGCAGGA | 36 |
| *PECAM1* | V | NM_000442.3 | TGGACTGTGTTGCTTTTCTTGACCACTTTGTCAATACCTGCAGTGCTTGAGCTCTAGGCCCAAAACGACCTTAATGGTCA | CGAAAGCCATGACCTCCGATCACTCGGGCATCATAAGAAATCCTGGGCTGGGAGAGCATTTCACATACGACTATC | 10 |
| *PENDRIN* | CD | NM_000441.1 | GATGGAAAATGATGCAGCCAGCATCTCCGAGAACAAGCTCACAGGTGGAACTTGGAGGAGTTGATAGTGGTAAAACAACATTAGC | CGAAAGCCATGACCTCCGATCACTCGTGGCATATACTTTTCCTACTGACACTGCAATAGCATAAGCCACCACAGC | 2 |
| *PIEZO1* | MS | NM_001142864.1 | GCGGCCGGCGAAGCTGACCACGATGAACATGCCAGCACACACATAGATCCTTCTGGAATTTCTTCCTTTGATTTTGCCATTTT | CGAAAGCCATGACCTCCGATCACTCGTGAGGCAGAGCAGGAAGAGGAACATGTAGACAATCTTGTAGACCACGAG | 37 |
| *PKD1* | CI | NM_001009944.2 | CGGAAGGTGTAAGAGATGGTAGGACCCCCAGGGATGGGCGCATCTCCATGACTGCTTGAGCGGCTGGAGAATCTG | CGAAAGCCATGACCTCCGATCACTCCTCGTTCTCAGCCGTGACGATGATATTGAAGGTGCCCACGGAG | 10 |
| *PKD2* | CI | NM_000297.3 | ATTCAACAGCACTGCATGCTCTTTTGCTTATTAGCGCTCAAAATGTCACCCTTTCGCCACCCATATAAACCCCACTTCGTCCTCA | CGAAAGCCATGACCTCCGATCACTCAAAATATAGCATTAAACGACAGGATTCTCTTGGTTCTCCAAGTGAGGGTT | 10 |
| *PKHD1* | CI | NM_138694.3 | ATAATATCAGGCTAACACCTTCCAAACTAGAGCCTCGGATGGTGGCCCAGCAAGGCAGAGCAAATGTGACACTGTCTATCAGTAC | CGAAAGCCATGACCTCCGATCACTCTCTGCTTGAATTGCTTGTAGCGACATTGATGGCACACGAGTAAGATCCAA | 10 |
| *PLAU* | IJ | NM_002658.2 | GAGCGACCCAGGTAGACGATGTAGTCCTCCTTCTTTGGGTAATCAATGAACAAAGCAAACCTAATGAAGCCAGACACGAGATCAC | CGAAAGCCATGACCTCCGATCACTCGGTTTTCCACCTCAAACTTCATCTCCCCTTGCGTGTTGGAGTTAAGCCTT | 38 |
| *PODXL* | PO | NM_005397.3 | CTCAATCAGATGAAACCTCACCATGCCCACTGCTTAGGAGCCTTCTACA | CGAAAGCCATGACCTCCGATCACTCACGATGGAGACCATGGCGA | 7 |

| | | | | | |
|---|---|---|---|---|---|
| | | | TCTAGCCCAGATCCTACGAGATGAGCTACGTAACTA | AAGTTCAACATTCCACACAGGATTCCCCCTT | |
| *POU3F3* | LH-DT-CD | NM_006236.1 | TTGGATCCCGGGCGGCGTCATGCGCTTCTCCTTTTCCATACGCATGACTACATTACAACGGGCCAGGAAG | CGAAAGCCATGACCTCCGATCACTCCCCACCTGCGAGTAGACGTCGTCGGGCGTCTGCTG | 2 |
| *RBFOX3* | N | NM_001082575.2 | TGGTGGCAGAATTTCCTCTGGCTGCTGTGACTTCAAGCTGCCTATTGAAGCAATCCTCTCCCCAATACTTAAAAA | CGAAAGCCATGACCTCCGATCACTCGGTTCCCAGCTGGAGGGGCTGCAGACCTGGGAAAA | 39 |
| *REN1* | SC | NM_016118.4 | GGAGGGAAAGTCTTGGATGCACCTGGCACTCAATCCACTCGGCCCAGCTGTCTCAATTATTGCTCAAAAACCACTATT | CGAAAGCCATGACCTCCGATCACTCCGCATGATCCATCCTGTCTTCAGTCAGTGCCTTCTGGAAG | 14 |
| *RET* | UB | NM_020975.4 | CTGGACGTTGATGCCACTGAATGCCTGGCAGTTTTCCACACAGACTTTCCCATCCTCTTCTTTTCTTGGTGTTGAGAAGATGCTC | CGAAAGCCATGACCTCCGATCACTCGTGACCACCCCTAGCGTGCTGCAGTTGGCACCAGAGGAATGCAGCTTGTA | 9 |
| *ROBO2* | PO | NM_001128929.2 | ACATACTTCACTTTGAAGCCTTGGTTTATTAATCACCAACACCTGCCAGTCTATCGTGGACCCAAAGTCGATACGTCCGATTAAG | CGAAAGCCATGACCTCCGATCACTCCAGGAAGGTATGATGCTATGCCCACGTTATTCCAAAAGTCATCTGCACAC | 5 |
| *SBSPON* | UB | NM_153225.3 | AAGTTCCTTGACACCGAGGATTACCAATTGCTTGCCAATGGAGAGTCTGACCGGGAATCGGCATTTCGCATTCTTAGGATCTAAA | CGAAAGCCATGACCTCCGATCACTCACTGTGAACAGCTGGACAAGAACACTGGTCTACTCGCCGAACTTTTTTCC | 12 |
| *SHROOM1* | UB | NM_133456.2 | CCTGAAAGTCCCAAGGCATCAGCTGGGACAGGTTTGAGGACCCGAAGCAATACTGTCGTCACTCTGTATGTCCGT | CGAAAGCCATGACCTCCGATCACTCGGCTAGGGCAGTATTGTGAGAGGGACCTGGAGTATCATTG | 12 |
| *SIX1* | MM | NM_005982.3 | GGCGGAGGAGTAGGGGAGGTTCTGGACACGGAACGGCTCCTAAGGTTGCTGATTTGGTTGTTGGAGACCCA | CGAAAGCCATGACCTCCGATCACTCAAGGCGAAAACCGGAGTCGGAACTTGGCGGTGGGCGGCCAAGGAAGAGAA | 3 |
| *SIX2* | MM | NM_016932.4 | AGGGAAAAGGCCTCAGGAGATGGCCTAGGTTCAAAGTCCCTGTATCAAGGCCTAGCCTAAAGGTTCTTGCAGAGCAACAT | CGAAAGCCATGACCTCCGATCACTCTTACTAGACGAAAAGCTGACACCTGCAGCTCCCACTGACCCACATGGGCG | 3 |
| *SLC12A1* | LH | NM_000338.2 | CCATATACAACAAATCCGATATGGATCCCTTTCTTGCCACGGGAAGGCTCCTCAGGTTGTTACTTGAAGGGTTCAACACGAGCTC | CGAAAGCCATGACCTCCGATCACTCTCTAACTAGTAAGACAGGTGGGAGGTTCTTTGTGAGGATTTCCAACCAAG | 4 |
| *SLC12A3* | DT | NM_001126107.1 | GTGGACAGGGATGTCAAAGCCTGGAGTTACTGGCAGTAAAAGGTGAGCACCATTAGCTCGGATGCTATCAGCTTGCGCCTATTAT | CGAAAGCCATGACCTCCGATCACTCCTGTGGACTAGAGGCAATCAAGTTCCAACTTATCCCACACACTCAGAGCT | 4 |
| *SLC22A3* | PT | NM_021977.2 | GACAGTGGAGGGCCTCTGGTTTGCTTTGAGAAGGACAAATACATTTTACACTTAGGCTACCAAATGAATTTAAAGCCAGCTGAAA | CGAAAGCCATGACCTCCGATCACTCGGCTGGCAGCCTAACCCTCACATGAATTTTTCCCTACCTCTATTTAGGGT | 4 |
| *SLC2A2* | PT | NM_000340.1 | AGCACCCTGCTAAGCTTTTGGGACCCATCAGAACTATATTTCCATAATTGCCTAATTAGCTCTAGGAAACACAACCCCGGGATTT | CGAAAGCCATGACCTCCGATCACTCAGGATTGATCAGTGCTCCAGTTGGTGGAGAAAACAGCCTAGAGATACGTT | 10 |
| *SLC34A1* | PT | NM_003052.4 | TCAGGAGCATCACGGCCACCATGGATGTTGAAGGAGGCTTTTTTGCAATTAAACTGCCCAGGCGATCTGTTG | CGAAAGCCATGACCTCCGATCACTCTGGATGATGAGCTTCGTGAAGGGCTCTGTGATGATCTTGAGCAGG | 10 |
| *SLC40A1* | PT | NM_014585.5 | ACAGAGATTATGGGCACAGATTCAGGACTTGTCTCCGGGACAATATTAGCCCAATGCTTGCAGTATGTATCCTGATCGTGCGTGC | CGAAAGCCATGACCTCCGATCACTCAGGACCAAAGACCGATTCTAGCAGCAATGACGCCTGCAAACAGCAGACTG | 4 |
| *SLC41A3* | DT | NM_017836.3 | ACCTTCTTGGCACAGCAGCAGTGTGGCCCACGGAGCACGATCTGTATTTTGCACCTTTCGCTATGCTGAG | CGAAAGCCATGACCTCCGATCACTCATGAAGGGTTGTATGAGGCCCCATCCTGGGGAGGCTGTAC | 4 |

| | | | | | |
|---|---|---|---|---|---|
| *SLC6A19* | PT | NM_001003841.2 | TGTAGGTCAGCTCCTGACTGACCTCTACCACGAAGAAGAAGAGGAAGATGCTTGAGTTATACGGAACTTCGCAAAAGTATTCCCT | CGAAAGCCATGACCTCCGATCACTCGGAGATCTTCTGGGATTTGGGAAATTCCTCGTAGCCAGGGTCCCAGATGC | 10 |
| *SLC8A1* | DT-UB | NM_001112800.1 | AAACGATGATGTGCTTTTTGAATGGGGTTATTCTTACTGTTTGCATAGGGCGTTCAATCGCTTAGTTTCGTGGCGGGATTTGAGG | CGAAAGCCATGACCTCCGATCACTCATCCCAAGCAACGGCAACATGGGACTGAGACACATGTTAATGCAAATGGA | 12 |
| *SLC9A3* | PT | NM_004174.2 | CGGTGACTGCGTCGTTCAGCAGCGACTCCCCGAAGACGATGATGAACAGGCCATCCATCAACAACTGCTCCAACAGCCTTTCCAT | CGAAAGCCATGACCTCCGATCACTCGTTGTCACCTCCCAGCGCCACGAAAGATTCAAACACATTGTACAGAACCA | 10 |
| *SNAI1* | EMT | NM_005985.2 | GATTGGGGTCGGAGGGCTTCCTGACGAGGAAAGAGCGCGGCATAGCATTACGACTTTACGAGTTCGCAGAACAAAGACTT | CGAAAGCCATGACCTCCGATCACTCTAAACTCTGGATTAGAGTCCTGCAGCTCGCTGTAGTTAGGCTTCC | 40 |
| *SOX2* | S | NM_003106.2 | ATCCTGCCGCCGCCGATGATTGTTATTATTATTTTTTGGAAAGGCTTAAGCCGATCTTCATAACGGACAAACTGAACGGGCCATT | CGAAAGCCATGACCTCCGATCACTCGGCAAACTGGAATCAGGATCAAAAAAAGCGCTTCCCTCCTCCTCTGGCCG | 29 |
| *SOX9* | UB | NM_000346.2 | TTTCTCGTTGATTTCGCTGCTCCATTTAGCCAAGGTTGGCCTGGCCACTGCACAATTCTGCGGGTTAGCAGGAAGGTTAGGGAAC | CGAAAGCCATGACCTCCGATCACTCATTCTCCATCATCCTCCACGCTTGCTCTGAAGAGGGTTTAAAAAGTCCAG | 9 |
| *SPP1* | LH-MX | NM_000582.2 | CTGACTCGTTTCATAACTGTCCTTCCCACGGCTGTCCCAATCAGAAGGCGCACTGGTTGGTTATCCAATGATCGACCTGGAGTCT | CGAAAGCCATGACCTCCGATCACTCTTGTGGCTGTGGGTTTCAGCACTCTGGTCATCCAG | 12 |
| *STC1* | CD | NM_003155.2 | GGATCTGGATACACTGAAAGCTTAGGTGAGGATTTGATGAGGATTTTTTTCTTTTAACAAACACTGGCTTTTTCCATCTTGTTGC | CGAAAGCCATGACCTCCGATCACTCTTCCTCTTTCCCTCTCCTGGCTTGAGTGAAGATGT | 41 |
| *SYNPO* | PO | NM_001109974.2 | ACAACAGTTGCCGGTGGGTTCTGATTGACATCGGTGGCTGCAAATGCACTCTATATGGAGGGAGAGTAGCTGGAT | CGAAAGCCATGACCTCCGATCACTCCTGAGGAATTCTGTTGGATGCTAGAAAGTGGCAGGCTCTGTGGG | 42 |
| *TACSTD2* | UB-CD | NM_002353.2 | AAGACATCCAAACTGCGTTCAGGCAGCTGAAACAGGCTTCTTTCCCAGTGCCGTCTCAGATGAGTGGGTTAATCAATCAAGTATG | CGAAAGCCATGACCTCCGATCACTCGGCAAGCTGAAGAATAAATAGACTGAGTTTCCGGGCAATGTCTGTCCTCA | 2,12 |
| *TBX18* | SC | NM_001080508.1 | CTACTGAAATTCCTCACCGTGTTATTTAACACCATACAGGCCTAGGAACTCTATGTCGTGCTATAATGGCGTCTGTCGTGCTCAT | CGAAAGCCATGACCTCCGATCACTCGCTTTGGCCTTTGCACTATAGCTCCCAAAAGACAGATTGCTGGAGATAAC | 6 |
| *TGFA* | DT | NM_003236.2 | CACAAAAGGCTGCACAGGTGATTACAGGCCAAGTAGGAAGGTCTGTGGCACCATCCACTTTCATGGAAACAATAAGAGCAGGGAA | CGAAAGCCATGACCTCCGATCACTCACACTGAATAACCCCAAGCAGACGGAGTTCTTGACAGAGTTTTGAAGGCC | 12 |
| *TGFB1I1* | SC | NM_001042454.1 | CCTTCTGCTCTCCTGAAGCCACCTTGCTAGATGGGAACTGAGACATGATTCTGGAATATCCTCATGCTCATAGATGTCAAATTTC | CGAAAGCCATGACCTCCGATCACTCGCTGGAAGGGAGGCTGGGTCTTTTCTTATCTTCAGACTGGTCCT | 43 |
| *TJP1* | PO-PT-LH-DT-CD | NM_175610.2 | GAGGCTTAAAACTGCCAGTGTCATTTACATCCTTCTTGGTCTCTAAGGATCATACGCCTTACGCCCCAAGGTTTAGCGTATCAAA | CGAAAGCCATGACCTCCGATCACTCGGGTTTGGGACCAATGATGGGAGCACCTGAAGGTTTAGATGCTACTTCTG | 44 |
| *TRPS1* | PT-UB-MET-MM | NM_001282902.1 | AAGCTATGTTCCTCCTTATGATTTAGTTCTGCAGCATCACTCTGATCTAGCTCAATTGCCATTGCTGGAATAGTGTCGGTA | CGAAAGCCATGACCTCCGATCACTCCTGCGCTTTTCAAGTCCTTCTTACTGCTAGAAGATGGATCTTGAACATGC | 10 |
| *TUBB* | HK | NM_178014.2 | ACTGCTGACACCTCCCTTGAAGCTGAGATGGGAAATGGACATACTTAGAACGCTCATTTTGAACATACGATTGCGATTACGGAAA | CGAAAGCCATGACCTCCGATCACTCACAGACTCCTCCAGAGTAGAGCTTGGAGGGAGATTGAAAGTGGAGATAAT | |

| | | | | | |
|---|---|---|---|---|---|
| *TWIST1* | IJ-EMT | NM_000474.3 | TCCTGGAAACGGTGCCGGTGCTGCAGAGCCCGCGACAGCGAATCTGCAACTAACGCAAGTTACATCCTAG | CGAAAGCCATGACCTCCGATCACTCAAAAAGAAAGCGCCCAACGGCTGGACGCACACCCCGCCAGGCC | 40 |
| *UMOD* | LH | NM_001008389.1 | TCATAGTTTCCAGCAAACCGGAACATCTGGACGGAAAATCGGCCCTGGGACGTCGTGTCTTAGACGACTGTGTGTGATTCTCGAG | CGAAAGCCATGACCTCCGATCACTCTTTCATTCATGGTGTCACAGAGATAGACTTCACAGTGCAGGTAGACTAGG | 8 |
| *UPK1A* | UR | NM_007000.3 | GCTCAAATGGCAATCACAAACGGTTCAGCAAGAATGGCGATGCCTGCCATCTCATGTCCTCTGTTAATCCAGCCTGAATATGCCA | CGAAAGCCATGACCTCCGATCACTCGACAGTAAAGTGACCGTGACAGAACTCTCATGGCAATAGAGGCTTCCAGA | 31 |
| *UPK1B* | UR | NM_006952.3 | TGCTTCAGGAAGAGGTTGGGTGTGAAAAAGTCTTGTTGTGTTGCTGCTGTCAAACTGGAGAGAGAAGTGAAGACGATTTAACCCA | CGAAAGCCATGACCTCCGATCACTCACTGGTCATCATTGTTTGGAGGGCTGTTGTTTTGGTACCTCTCTAGCATC | 31 |
| *UPK2* | UR | NM_006760.3 | ACGAGGTTTGTCACCTGGTATGCACTGAGCCGAGTGACTGCGATTGCTGCATTCCGCTCAACGCTTGAGGAAGTA | CGAAAGCCATGACCTCCGATCACTCTCCCCTTCTTCACTAGGTAGGAAATGTAGAATTTGGTTCCTGGC | 31 |
| *UPK3* | UR | NM_001167574.1 | GGGCCTCTTTGCTGTCAAACATGCAGAGAGGCTTTTCGCCCTGGAACAACGGTTATATTCTCTGGCAGAAC | CGAAAGCCATGACCTCCGATCACTCCAGGACATACAGGTAGACCTCGTGGGTGCCAGTGA | 31 |
| *VEGFC* | DT | NM_005429.4 | ATTTTCCAATATTCTGGGTAGAGTACAGTCATGAGTTCATCTACACTGGACCGAGTGCATGAGCTGTCTTTCACATGATACATCG | CGAAAGCCATGACCTCCGATCACTCCCTGTTCTCTGTTATGTTGCCAGCCTCCTTTCCTTAGCTGACACTTGTAC | 12 |
| *VIM* | EMT | NM_003380.3 | CGAATGTGCGGACTTTTTGTTAGCAAGAAACTTCTGCAGCCTTTGGAGAGCTGTGACATCGCTTCTAGAGTATGAGCCGCAATGC | CGAAAGCCATGACCTCCGATCACTCGACAAGAGCGCCCCTAAGTTTTTAATAACTCGCTAAAGCCTGTCTTTGCT | 40 |
| *WNT11* | UB | XM_011545241.2 | CAAAGCACAAGCCGCCTCCCATGCCTGGCATCTGGAATTCACAAGCCTGGAGTTTATGTATTGCCAACGAGTTTGTCTTT | CGAAAGCCATGACCTCCGATCACTCCAGACCTTCTGTTCCTGGTGGCTTCCAAGTGAAGG | 9 |
| *WNT7B* | UB | NM_058238.1 | CAGCCTGGGAACTGGTCTGGAGAGGCCAGCCGGCGTAGCTTTTCTGTGTCCTGTTGAGATTATTGAGCTTCATCATGACCAGAAG | CGAAAGCCATGACCTCCGATCACTCTAATTTGTGTCAACATCTGTCCCCACCGCCCCAGGGCCCAGCGGCAGACC | 18 |
| *WNT9b* | UB-CD-MET | NM_003396.1 | CTTGGTCTCCCTTATTTCCCTCCATGTCTTTCTCACTCTGCCAAAGACGCCTATCTTCCAGTTTGATCGGGAAACT | CGAAAGCCATGACCTCCGATCACTCCTCTCTCCCGTCCAAAATTTCAGTCCTGCTGTGCT | 18 |
| *WT1* | PO | NM_000378.3 | TGACCTCGGGAATGTTAGACAAGATTCACCCCCGATGCCTTGCTCTCTGACGCTCACGTGATCTACCCTAGCTGACCGCTAATGA | CGAAAGCCATGACCTCCGATCACTCAAAAGTTGCCTGGCAGAACTACATCCTGCTTTCCAGGTTAGCAGCCTGGC | 42C |

### Supplemental Table References:

11. Paunescu, T. G., Da Silva, N., Marshansky, V., McKee, M. & Brown, D. Expression of the 56-kDa B2 subunit isoform of the vacuolar H+-ATPase in proton-secreting cells of the kidney and epididymis. *American Journal of Physiology-Cell Physiology* 287, C149–C162 (2004).

33. Tan, R. J. & Liu, Y. Matrix metalloproteinases in kidney homeostasis and diseases. *American Journal of Physiology - Renal Physiology* 302, (2012).
